# Generalization and Transfer of Navigation Memory in Honeybees

**DOI:** 10.1101/2022.10.01.510441

**Authors:** Eric Bullinger, Uwe Greggers, Randolf Menzel

## Abstract

Flying insects like the honeybee learn multiple features of the environment for efficient navigation. Here we ask whether the memory of such features is generalized to novel test conditions. Foraging bees from colonies located in 5 different home areas were tested in a common area for their search flights. Harmonic radar was used to track the search flights of the bees. The test area resembled partly or not at all the layout of landmarks in the respective home areas. Random search as the only guide for searching was excluded by two model calculations. The generalization and transfer effect was quantified by a partial least squares regression analysis of the guiding effect of the ground structures in the test area. Classification was performed with a support vector machine in order to account for optimal hyperplanes. A rank order of well separated clusters was found that partly resemble the graded differences between the ground structures of the home areas and the test area. We conclude that foragers guide their search flights by a generalization and transfer process that is based on their acquired navigation memory. The conclusion is discussed in the context of the learning and generalization process in an insect, the honeybee.

**Summary Statement:** A new test procedure is applied to address the question whether a flying insect, the honeybee, uses a navigation memory that can be generalized to novel environmental conditions.

## 1 Introduction

Honeybees explore the environment around their hive and form a memory that brings them back to their colony. Like in other animals, the structure of this memory is still under debate (M. Collett, Harland, and T. S. Collett, 2002; Jacobs and Menzel, 2014; Wiener et al., 2011). Initially, it was proposed that bees navigate according to an egocentric reference frame only because the vanishing bearings of bees released at an unexpected site indicated a flight vector the animals would have taken if they were not translocated (Dyer, Berry, and Richard, 1993; Menzel, Geiger, Chittka, et al., 1996; Menzel, Geiger, Joerges, et al., 1998; Wehner and Menzel, 1990). However, bees return home rather quickly from all directions around the hive within distances of < 500 m to the hive (Menzel, Brandt, et al., 2000). The time of returning home may differ depending on the structure of the landscape (Dyer, 1996), but they are usually not lost. Additional insight came from data which were collected using harmonic radar to track their flight paths after release (Menzel, Greggers, et al., 2005; Menzel, Kirbach, et al., 2011). After bees followed the egocentric vector component they returned home directly or steered to the feeding site first before returning home. One possibility of explaining these results is to assume that bees refer to picture memories of the landmarks very much as they do when aiming into the hive or a feeding place from the immediate surrounding (Cartwright and T. S. Collett, 1983; T. S. Collett, Ibarra, et al., 2013; Stürzl et al., 2016). However, the conditions during homing flights over distances of many hundred meters differ from aiming in to the nest or feeding place over a few meters because the patterns of large scale landmark features will be effected by partial or complete occlusions, distance dependencies and sequences of landmarks. Notably, elongated ground structures may be particularly salient and informative in this context (Brebner et al., 2021; Menzel, Tison, et al., 2019) because they are experienced as such only by extended flight, they provide learned compass direction (Dyer and Gould, 1983; Frisch and Lindauer, 1954), they may be polarized in relation to the intended goal when combined with compass information and they could uniquely specify locations when combined with other landscape features. Still, bees may apply stepwise match-miss match strategies for further distant landmarks as suggested by Cartwright and T. S. Collett, 1983. Thus the question arises whether the navigation memory is a concise memory or is composed of multiple independent functions of guidance.

Large scale landscape memory acquired during orientation flights and foraging flights has been characterized so far mostly by translocating animals to other sites within the explored area and monitoring their flights back to the hive. Here we apply a different approach. Animals from a colony located at a different site (its home area) are transported into a test area that partially resembles landscape features on the ground but differs drastically with respect to rising objects both close by and at the horizon. Five different home areas were chosen, and all animals were released at the same place in the test area. Four home areas were so far away from the test area that no test bees ever came close to the test area. One home area was located 1.6 km away from the test area, and indeed a few of these animals managed to fly back to their home area. These few animals were not included in our analyses.

All animals were experienced foragers having calibrated their sky compass and visual odometer, and learned the multiple landscape features for successful homing. It is known that foragers transported into an unknown area return multiple times to the release site (Dyer, 1996; Menzel, Geiger, Joerges, et al., 1998), possibly applying irregular search flights. Thus, we expect that the animals will perform random search flights with multiple returns to the release site if they do not recognize any even partially matching guiding features. If, however, their navigation memory can at least be partially generalized to the features experienced in the test area, their behavior will differ from random search. Generalization between home area specific memories and experienced features in the test area may motivate them to explore these features more intensively. The local density of exploration may thus reflect a generalization and transfer effect that may support the notion of a concise navigation memory.

The navigation memory established in the 5 home areas differs due to the layout of the respective landmarks, potentially resulting in different search patterns in the same test area. Thus, a similarity measure based on the search patterns may reflect components of the navigation memory. We hypothesize that this memory transfer indicates a form of navigation memory based predominantly on salient elongated landmarks. Matching of stored views of the panorama with views in the test area should play no or little role in our experiments because of the drastic difference of panorama between the home areas and the test area for four of the five home areas. The same will apply to localized rising landmarks because there were none such landmarks in the test area. Ground structures, however, may influence their search flights. In the test area, the animals may thus identify features preferentially on the ground that partially resemble features they had learned in their home area, and thus they may generalize between such features.

First, we shall examine whether the search flights follow a random search strategy running two mathematical models. After showing that random flight alone cannot explain the search behavior, we find that the search flights differ between animals from the different home areas. We next asked about the impact of the elongated ground structures in the test area. Finally, we quantify the differences in search strategy of animals from the different home areas and compare the effects by analyzing the differences between the elongated ground structures of the home areas with that of the test area.

## 2 Methods

The experiments were carried out close to the village Klein Lüben (Brandenburg, Germany). The area is characterized by open grass land, agricultural land, forest, creeks, irrigation channels, small roads and the river Elbe. The harmonic radar for flight tracking in the test area was placed at the coordinates 52°58′31.14″N 11°50′11.35″E. The test bees were released at the Site R (Fig. 1F, 52°58′36.37″N 11°50′13.37″E). Five colonies of *Apis mellifera carnica* were positioned in five different areas (home sites A to E, Fig. 1) that differed with respect to the respective landscape structures. Hive A was located at 52°59′07.94″N 11°49′05.34″E, 1.64 km NW of Release Site R, and Hive B at 52°58′53.18″N 11°48′30.87″E, 2.1 km E of R. Both areas consisted of predominantly agricultural fields, a road, and irrigation channels. Hive A was close to a row of high rising poplar trees, and Hive B close to isolated low trees. The home area around Hive C (52°57′22.23″N 11°51′42.29″E, 2.89 km SW of R) was characterized predominantly by open meadows with irrigation channels and an edge of a small forest in the N. Hive D (53°00′08.98″N 11°52′17.75″E, 2.21 km NE of R) was located in the middle of a large forest with a SE–NW stretching forest aisle. Hive E (52°58′46.02″N 11°46′15.86″E) was placed at the bank of the river Elbe overlooking a country side with scattered trees. The distance between Hive E and the radar location was 4.55 km. The river was between 350 m and 450m wide during the years 2009 and 2010. In 2011 the river flooding lead to a width of >1000 m of open water. Figure 1 gives an impression about the similarity/difference of the elongated ground structures in the test area and the 5 home areas. Notice that the layout of the ground structures in the test area was more similar to that in home areas A and B than to C–F and that home areas E and F were very different between each other and the test area.

**Figure 1:**
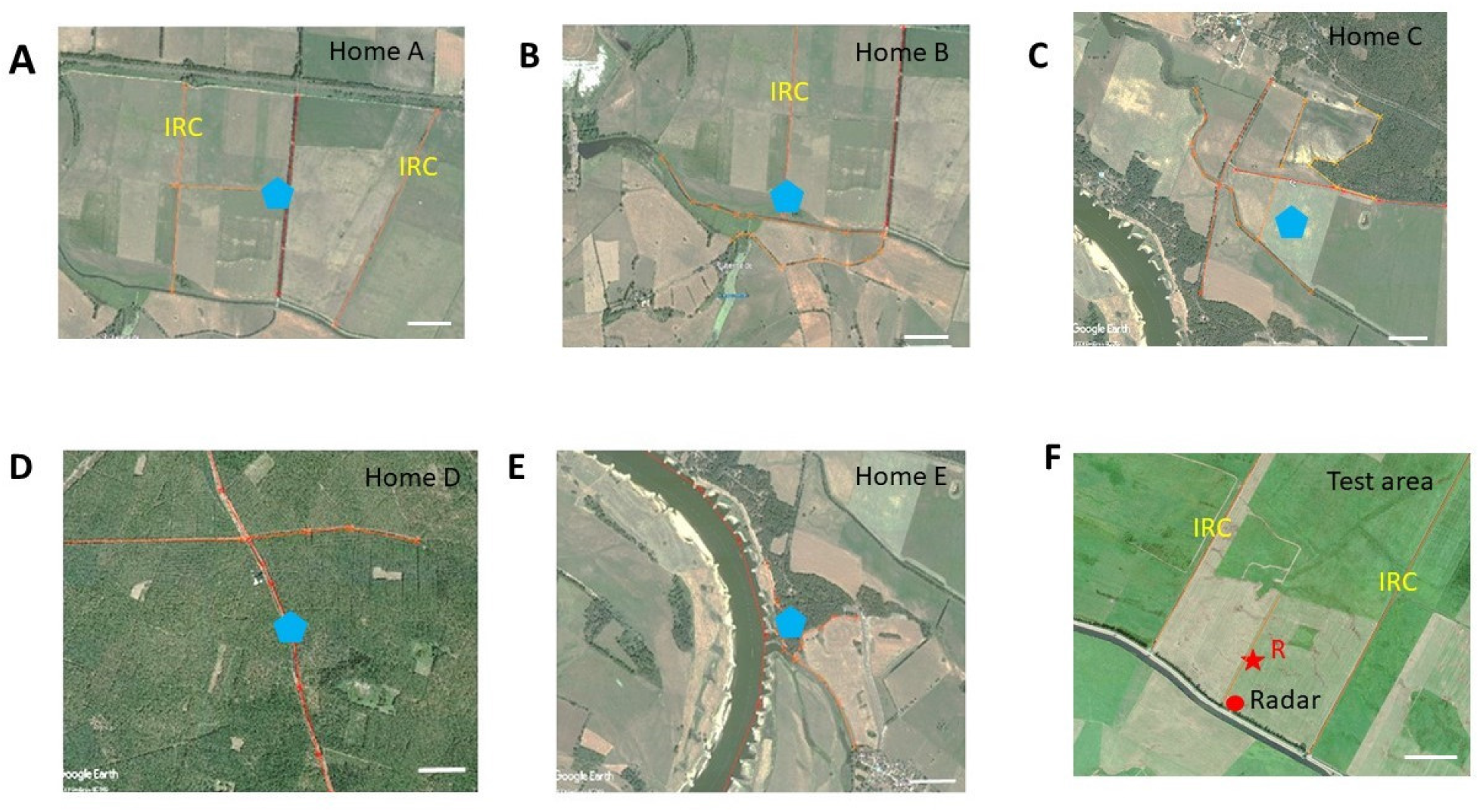
Areal views of the test area (F) and the 5 home areas. **(A-E)** The release site R in the test area is marked with a red star, and the location of the radar with a red dot. The locations of the hives in the five home areas are marked with blue pentagons. The elongated ground structures in the test and home areas are highlighted with orange lines. The scale in each subfigure corresponds to 200 m. IRC: irrigation channels. Notice that the ground structures in the test area are more similar to home areas A and B than to C–E, and that home areas D and E are highly different from the test area and from each other.

The experiments were carried out in the summers (July to September) of the years 2009, 2010 and 2011. The colonies were placed at the respective sites at least 4 weeks before the experiments started, ensuring that only foragers were tested that were familiar with the home area and had not experienced any other area. Foragers of each colony were trained to a feeding site very close to the respective hive (< 7 m). Single foragers were then collected at the feeder after they had sucked their fill, transported in a dark box to the test area within < 30 min, equipped with a transponder for tracking their flight with harmonic radar and released at the release site R in the test area (Fig. 1). All test bees from the 5 colonies were released at the same respective site R. The test area was characterized by two irrigation channels (IRC) NW and NE of the release site R. They ran parallel to each other at a distance of 611 m stretching from SW to NE. In addition, a borderline between two grasslands mown at different times ran parallel to the IRC, roughly in the middle of the large and otherwise rather evenly structured grassland. Release site R was located close to this borderline. Additionally, several rather weak ground structures characterized the test area: lines in the grass running parallel to the borderline originated from mowing of grass, patches of grass growing at different height, and a dip in the ground running through most of the test area from SW to NE contrasting grassland with different grass species. The test area’s border running SW–SE was characterized by a row of bushes (at the closest 30 m SE of the radar station) running parallel to a small road and a creek. There were no trees or other high rising structures within 1 km of the radar other than the radar station itself (height 6m) and the row of bushes behind the radar. The home areas of the 5 hives were selected to either resemble partially the elongated ground structures in the test area (e.g. the two IRC, the border between two grasslands and the row of bushes) or to be different in various respect (rising close or/and far landmarks) from the landscape structure of the test area. The search flights of the bees were tracked from the radar fixes collected every 3 seconds with a customs made program that converted the fixes from the circular display to a Cartesian map (Cheeseman et al., 2012). Finally, the fixes were imported into Matlab and overlaid for visualization purposes to test site’s Google map. The map was also used to quantify the landscape features in the test area. The analyses of the flight tracks included a statistical evaluation of their spatial relations to landscape features of the respective home site. The procedures will be explained in the Result section.

The release site in the test area was chosen such that most of the search behavior could be monitored as indicated by the finding that during most of a test flights the respective bee was seen on the radar screen. Between 30% and 50% of the bees (depending on home area) left the scanned area for short periods. Most of them returned to the release site. No information was available where the bees from the different home areas had foraged. No spots of dense forage (e.g. flowering tree or bush) were found within the natural range (up to 2 km radius) of the respective hives during the experimental periods (July, August, September). We, therefore, assume that bees foraged on widely scattered flowers, and thus the individual bees from the different hives may have foraged in different directions and over different areas from their respective hive. The colonies were regularly inspected and ample food store (pollen and nectar) was found in all cases. The test area was the most even and spacious pastry we could find (and where the farmers allowed us to work). The flat and horizontal level lacked any obstacle besides the row of bushes, an ideal condition for harmonic radar tracking. The skyline was flat within ≤ 2° visual angle around the release site over a large proportion of the area within which the test bees performed their search flights (Menzel, Greggers, et al., 2005). This applies to all of the area scanned by the radar besides the area up to 200 m NE from the row of bushes behind the radar.

We tested a total number of 50 bees leading to one flight trajectory each. Each animal was released only once, because no animal could be recaptured. The numbers of tested animals (N) and the total number of radar fixes (n) were for the different colonies: home area A N = 14, n = 6098, home area B N = 11, n = 4128, home area C N = 5, n = 1722, home area D N=13, n=4611, home area E N = 7, n = 2496.

### Radar tracking

Tracking bees with a harmonic radar was achieved as described in Cheeseman et al., 2012. We used a system with a sending unit consisting of a 9.4 GHz radar transceiver (Raytheon Marine GmbH, Kiel, NSC 2525/7 XU) combined with a parabolic antenna providing approximately 44 dBi. The transponder fixed to the thorax of the bee consisted of a dipole antenna with a Low Barrier Schottky Diode HSCH-5340 of centered inductivity. The second harmonic component of the signal (18.8 GHz) was the target for the radar. The receiving unit consisted of an 18.8 GHz parabolic antenna, with a low-noise pre-amplifier directly coupled to a mixer (18.8 GHz oscillator), and a downstream amplifier with a 90MHz ZF-Filter. A 60 MHz ZF-Signal was used for signal recognition. The transponder had a weight of 10.5 mg and a length of 12 mm. We used a silver or gold wire with a diameter of 0.3 mm and a loop inductance of 1.3 nH. The range of the harmonic radar was 900 m. Several limitations of the applied methods and the test conditions need to be mentioned. The range of the radar was limited to 900 m radius and did not scan through an angle of 150° to the SW due to the switch-off of the radar beam (radar blanking) and a row of bushes along a small road. Therefore, data about the searching behavior were available only within a limited sector (210°).

### Random Bees

Based on a modified version of the ant model by Wehner and Srinivasan, 1981, we developed a model of an artificial bee that explores the surrounding of its release site with a flight pattern that is independent of the landscape. Starting at the release in a random direction, the “bees” fly 10.000 steps of unit length with heading changes from step to step as specified by random numbers generated as the arctan of a normal distribution (zero mean, standard deviation 0.25) and filtered by a tenth-order low-pass digital Butterworth filter with normalized cut-off frequency 0.05. Whenever the distance to the release site is larger than 1 + k/10 unit lengths, with k = 1, 2, … the index of the flight loop, the heading angle is rotated by an extra 180°. This rotation is also added when the simulated bees comes closer than 0.1 unit length to the release site. This raw path is then smoothed by a moving average filter of length 25. Finally, comparing 1000 simulated paths to those of all real bee, the unit length was fitted to 2.22 m to achieve the same median distance from the release site as in the real flight data. The model bees are labeled S when their path include the sector not covered by the radar (radar blanking), and labeled R when only considering the circle segment covered by the radar, see Fig. 2C for an example.

**Figure 2:**
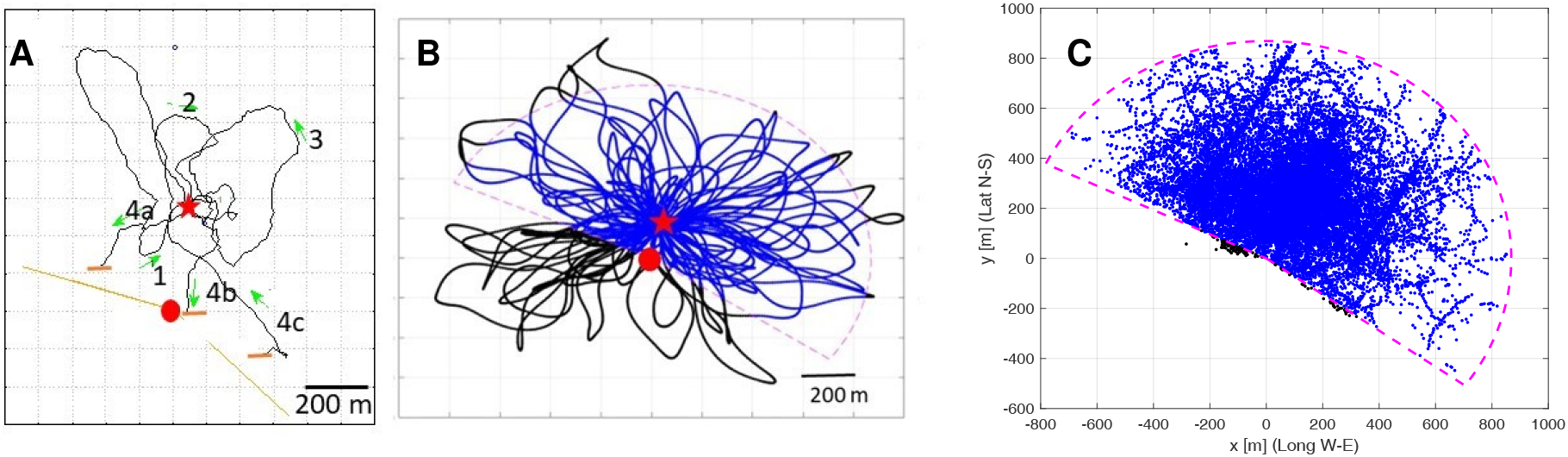
The structure of search flights. **A**: Example of a bee’s search flights. **B**: Example of a simulated bee’s flight. Two models were run, Model S with search loops in all directions (black and blue trajectories) and Model R in which fixes outside the radar range (dashed line) were excluded (only blue parts of the trajectories). **C**: All fixes of all search flights plotted together with the radar range with the radar location at the origin. Fixes in the range in blue, in black the approx. 0.45% outside that range.

1000 random bees were simulated for quantifying the scaling, but in the analyses, only 16 random bees (with and without radar blanking) were used to have a similar number as test bees per group.

### Computations

The calculations were performed, except noted otherwise, with Matlab (R2020b) and its Statistics and Machine Learning as well as the Image Processing toolboxes. The radar fixes were available to Matlab in radar-based Cartesian coordinates. A linear transformation to release-site coordinates, Cartesian as well as polar coordinates, was then applied.

For the circle statistics, the data points were transformed into polar coordinates around the release site. Multimodal non-parametric likelihood-based model calculations were performed on the angular data using R version 4.0.3 with the library `CircMLE` (Fitak and Johnsen, 2017) and the Rao test library by Landler, Ruxton, and Malkemper, 2019. For visualization, the analysis results were imported into Matlab. The transfer between Matlab and R was achieved via writing and reading appropriate text files. `CircMLE` fits a van Mises distribution (Mardia, 1972; Mises, 1918) to unimodal data u(*θ*). A van Mises probability density function has the form

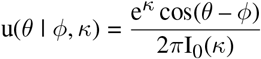

where I_0_(*κ*) is a modified Bessel function of order zero, *φ* the principal direction of the distribution, *κ* its concentration parameter (small *κ* for a distribution close to uniform, large *κ* in case the data clusters around *φ*) and –*π* ≤ *θ* ≤ *π* the angle. `CircMLE` fits bimodal data with bimodal distributions b(*θ*) that are a convex combination of two van Mises distributions, i.e.

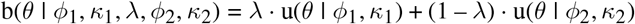

with 0 < λ <1 (Fitak and Johnsen, 2017; Schnute and Groot, 1992).

For determining significant differences between pairwise groups, we applied Kruskal-Wallis significance tests (Kruskal and Wallis, 1952) and the Measures of Effect Size (mes) based on Cohen’s U3 test (Cohen, 1988), the latter using the toolbox “Measures of Effect Size” by Hentschke and Stüttgen, Version 1.6.1 (Hentschke and Stüttgen, 2011).

For the heat map and edge analyses, the fixes were resampled by linear interpolation to obtain ten additional points equally distanced between fixes. The heat maps were obtain by first smooting the resampled data points using a 2D moving average filter of kernel size 50 m × 50 m, thus smoothing them up to a chessboard distance of 50 m. For the edge analysis, the Euclidean distance from each landscape feature to each of the resampled fixes was calculated in order to determine the closest landscape feature, the distance to it as well as the angle between the closest landscape feature and the flight direction. These data was then statistically analyzed. The partial least square regression was calculated by Matlab’s `plsregress`, the minimum volume ellipsoids were obtained by `MinVolEllipse` (on Matlab Central, by Nima Moshtagh, 2009). For obtaining the separating hyperplanes, called separatrices, these ellipsoids were resampled with 441 points, the optimal separatrix was calculated using Matlab’s `fitcsvm` with linear kernel functions, which is using a support vector machine approach. The distance between ellipsoids was obtained with the help of the Ellipsoid Toolbox (Kurzhanskiy and Varaiya, 2006), which uses YALMIP (Löfberg, 2004). Further functions from Matlab Central: `maxdistcolor` by Stephen Cobeldick (2020), `distinguishable_colors` by Timothy E. Holy (2011), `plotBarStackGroups` by Evan Bollig (2011).

Boxplots are used to illustrate distributions, and are particularly useful for non-Gaussian ones. The red line depicts the median, the second quartile, while the blue box goes from the first to the third quartile, ie. covers half the data. The box height is called inter quartile range, IQR. The notch on each box, extending 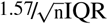 above and below the median, with n the number of data points, gives the 95% confidence interval for the median (McGill, Tukey, and Larsen, 1978). The lines up and down, called whiskers, extend from the box up to the last data points that is not further than 1.5 IQR from the first or the third quartile. The data points further away are outliers and marked by a plus sign.

## 3 Results

### 3.1 Random vs non-random searches

The basic search strategy of all animals released at the unexpected and unknown site in the test area consisted of multiple returns to the release site via multiple loops ranging over different distances and in different directions (Fig. 2A). No systematic sequences of growing distances and changing directions were apparent. One may assume, therefore, that the bees just performed random search flights. We tested this question by running two models of random search. Our model calculations assumed multiple returns to the release site with randomly directed loops of increasing size (Fig. 2B). These search paths were generated based on a modified version of the ant model by Wehner and Srinivasan, 1981 (see Section Methods for the details). The S model bees include the sector not covered by the radar (radar blanking), while the R bees paths are identical to the S bees, but excluding the fixes in the radar blanking (Fig. 2B). Figure 2C shows all fixes of all real bees together with the assumed sector covered by the radar that captures over 99.5% of the fixes.

If bees from the 5 different home areas would apply a random search strategy only, they should explore the area around the release site about equally frequently and no differences would be expected from the model R or S simulated bees. In a first step we compared the relative number of fixes in 16 angular sectors around the release site by normalizing it in each sector to that of the simulated bees of model R on the level of the individual bees from the 5 home areas and those of the modeled “bees” in model S. We chose the results of model R because it takes into account the bias of no fixes in the area not scanned by the radar, the radar blanking sector. Fig. 3E shows the directions from the release site to the respective fixes for 9 different distances. The relative distributions of test bees from all home areas taken together for all distances were statistically analyzed using two different procedures, the Kruskal-Wallis test and the Measure of Effect Size (mes) based on Cohen’s U3 test for two samples (Cohen, 1988) as implemented by Hentschke and Stüttgen, 2011 (Fig. 3A to E, File S1). Classical significance tests, e.g. null hypothesis significance testing, depend on sample size, while effect size does not. This mes compares two populations, A and B, and returns the percentage of population A that lies above the median of population B. Thus, the mes always lies in the interval [0, 1] as 100% = 1, and mes = 0.5 corresponds to equal medians. The further away a mes is from 0.5, the more significant the two populations differ. This difference is denoted by Δ_mes_ where Δ_mes_ = |mes – 0.5|. According to Hentschke and Stüttgen (2011), mes is able to uncover important aspects of the data that standard null hypothesis significance testing does not make visible.

**Figure 3:**
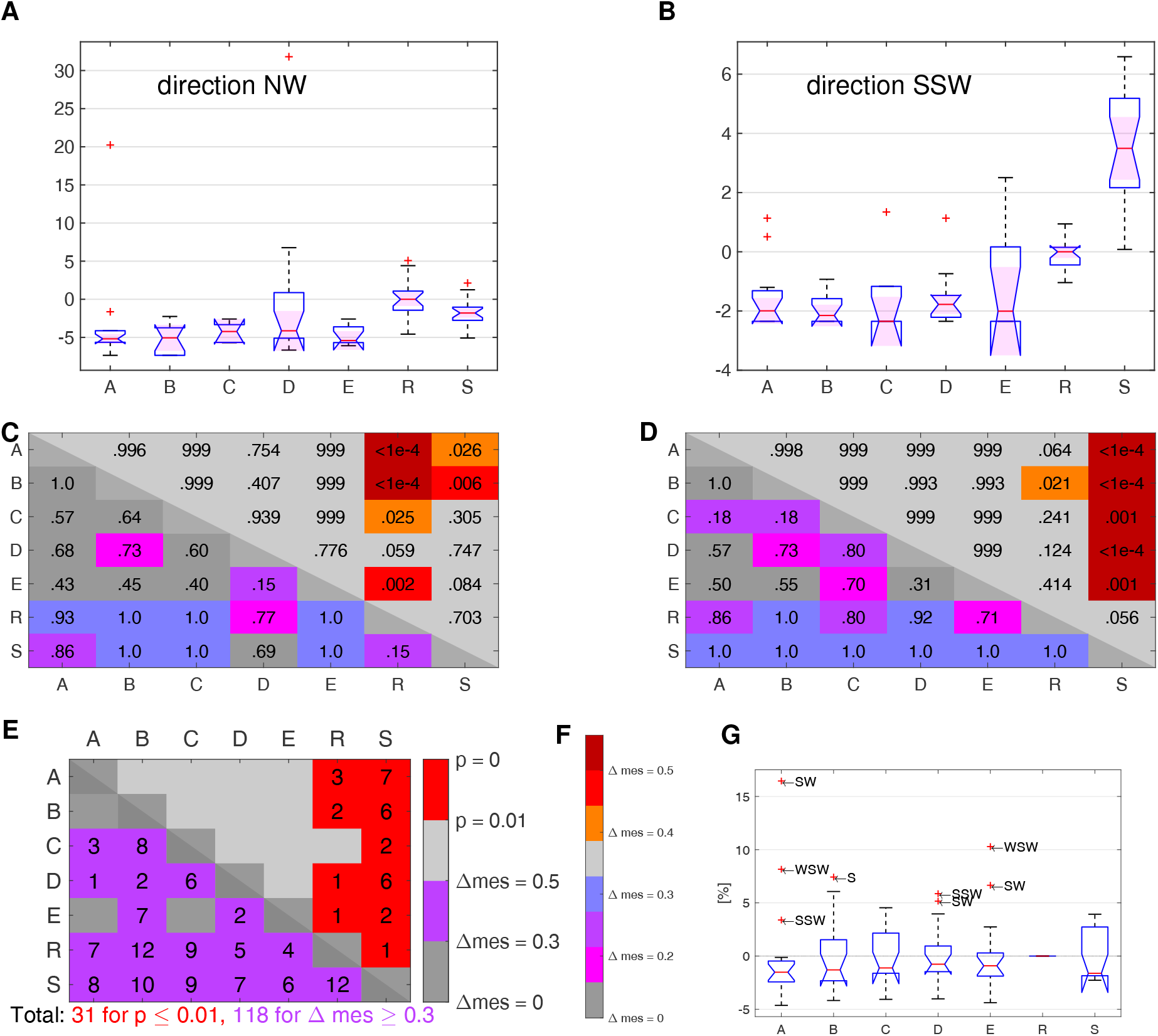
Relative proportions of flight directions for the bees from the five home areas (A–E) and the simulated bees in the models R and S. **A**–**E**: Statistical analyses of the frequencies of directions. The 16 directional sectors were tested separately. Two examples are shown here, with the full set of results in File S1. **A**&**C**: Sector NW, **B**&**D**: Sector SSW.**A**&**B** show the relative frequencies of the directions (in percent, additive normalization to the median of model R) for the 5 test groups and 16 “bees” each of the two models R and S. **C**&**D** mark the results of the two statistical analyses applied, the Kruskal-Wallis test (upper right side, yellow to red marks), and the Measures of Effect Size based on Cohen’s U3 test for two samples (see text). The color code the significance levels as depicted in **F**. **E**: Total number of significant differences for the two statistical calculations and the total number of significant cases for p ≤0.01 and Δmes = 0.2, respectively.

Significant differences according to the measures of Effect Size as expressed by Δmes were calculated for three values ≥ 0.1, 0.2, and 0.3, and these values are given in Fig. 3A to D for two examples (NW and SSW) of the 16 directional sectors together with the relative frequencies per bee of the directions for the 5 groups of test bees and the “bees” of the two models, R and S. The full set of the results of all 16 directional sectors are given in File S1. Although the radar blanking implies a dominance of flights into the N to E sector, several bee groups spent more time in south-western direction as compared to the R bees (Fig. 3G). The bees from the different home areas differed in their relative directional distributions both in comparison with that of the models and between the 5 test groups as indicated by the pairwise measure of effect size (Fig. 3D). The Kruskal-Wallis test is less selective and indicates significant differences only between the two model calculations and the 5 groups of test bees (total number: 27). The respective numbers of statistically significant cases for the measures of effect size are much more frequent with a total number of 183 including the two models, and 31 between the 5 groups of test bees only (Fig. 3D), where significant means p ≤ 0.01 or Δmes ≥ 0.2. While the Kruskal-Wallis test is only able to show pairwise significant differences between any test bees and model best, except for C vs. R, the measures of effect size shows significant differences between 20 out of 21 pairs, the exception here is C vs. E (Fig. 3E).

### 3.2 Non-parametric likelihood-based models of directions

Statistical analyses of circular data are usually based on null-hypotheses that assume unimodal distributions (Ruxton,2017; Steel et al., 2013). This might not be the case with our data. Since axial distributions (opposing directions) of bimodal distributions can be excluded (Fig. 3A), circular statistical procedures suitable for unimodal distributions, e.g. the Rayleigh test (Batschelet, 1981) cannot be applied. We, therefore, applied multimodal non-parametric likelihood-based model calculations (Fitak and Johnsen, 2017) to compare the distributions of directions of fixes as seen from the release site (angle *β*). These calculations are based on von Mises distributions (circular normal distributions) and test simultaneously null and multiple alternative hypotheses. The results of the model calculations for the distributions of *β* for the various groups were then evaluated on the basis of the Akaike information criterion (AIC), (Akaike, 1973). The AIC was applied to select the optimal model as a measure of uni- or bimodality with the smallest AIC as the best model. Following Fitak and Johnsen, 2017, an AIC relative to the best model with ΔAIC < 2 was considered to be well supported. Ten different types of models were proposed by Schnute and Groot, 1992. We determined for each bee the best models and present the respective parameters (*φ*_1_, *κ*_1_, λ, *φ*_2_, *κ*_2_) (Fig. 4). Bimodal distributions are described by models M3-M5 and unimodal distributions by models M2. The two examples shown in Figure 4 belong to one of the R model bees (Fig. 4A1–4) and to one bee from home area A (Fig. 4B1–4). All other results of this analysis are presented in File S2. Table 1 summarizes the result by listing all best models for each animal.

**Figure 4:**
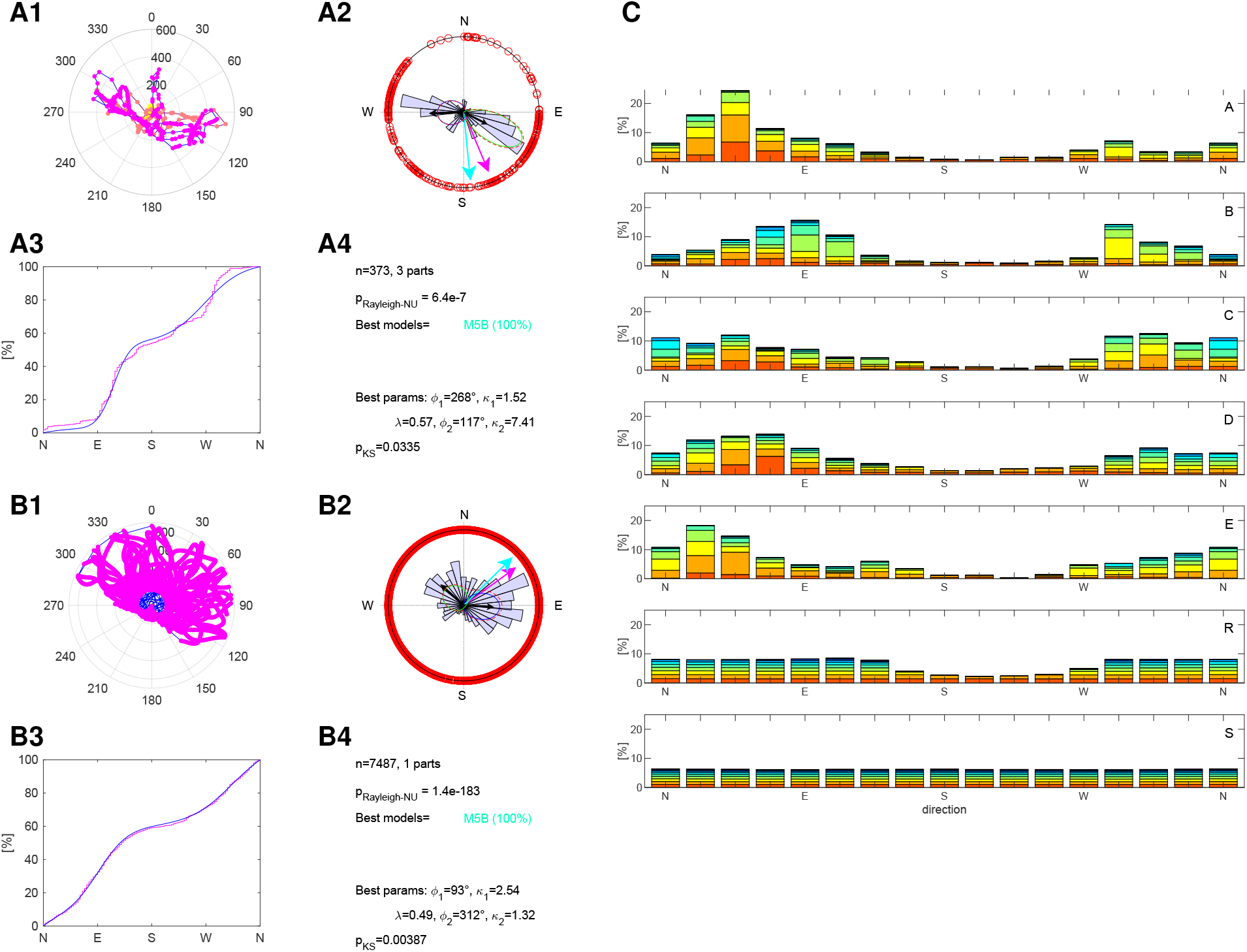
Angular statistics and path. **A1**–**A4** show the analyses for one simulated bee of model R, and **B1**–**B4** those of Bee 01 from Area A.File S2 shows the analyses for all simulated and tested bees. **A1** and **B1** depict an overlay of the flight trajectories. **A2** and **B2** are the corresponding angular distributions of the directions as seen from the release site. The black arrows mark the principle directions of the von Misses distributions (*φ*_1_ and *φ*_2_) together with the averages of the overall distributions (magenta arrow: median, cyan arrow: mean). **A3** and **B3** show the cumulative angular distributions for the best model (blue line) together with the measured distributions (red line). **A4** and **B4** summaries the most relevant values resulting from the multimodal non-parametric likelihood-based model calculations (*φ*_1_, *κ*_1_, λ, *φ*_2_, *κ*_2_). p_Rayleigh_NU_ gives the p-value for the Rayleigh test of non-uniformity. Random circular distributions can be rejected for small p-values. p_KS_ gives the p-value of the Kolmogorow-Smirnow test. The result of the model calculations can be rejected for large p-values. **C**: The directions from the release site to the respective fixes are given for 9 different distances depicted in different colors as indicated by the right bar. The bias in the frequencies across the 16 angular sectors in the random model R results from then cutoff of fixes in the not scanned area.

**Table 1:**
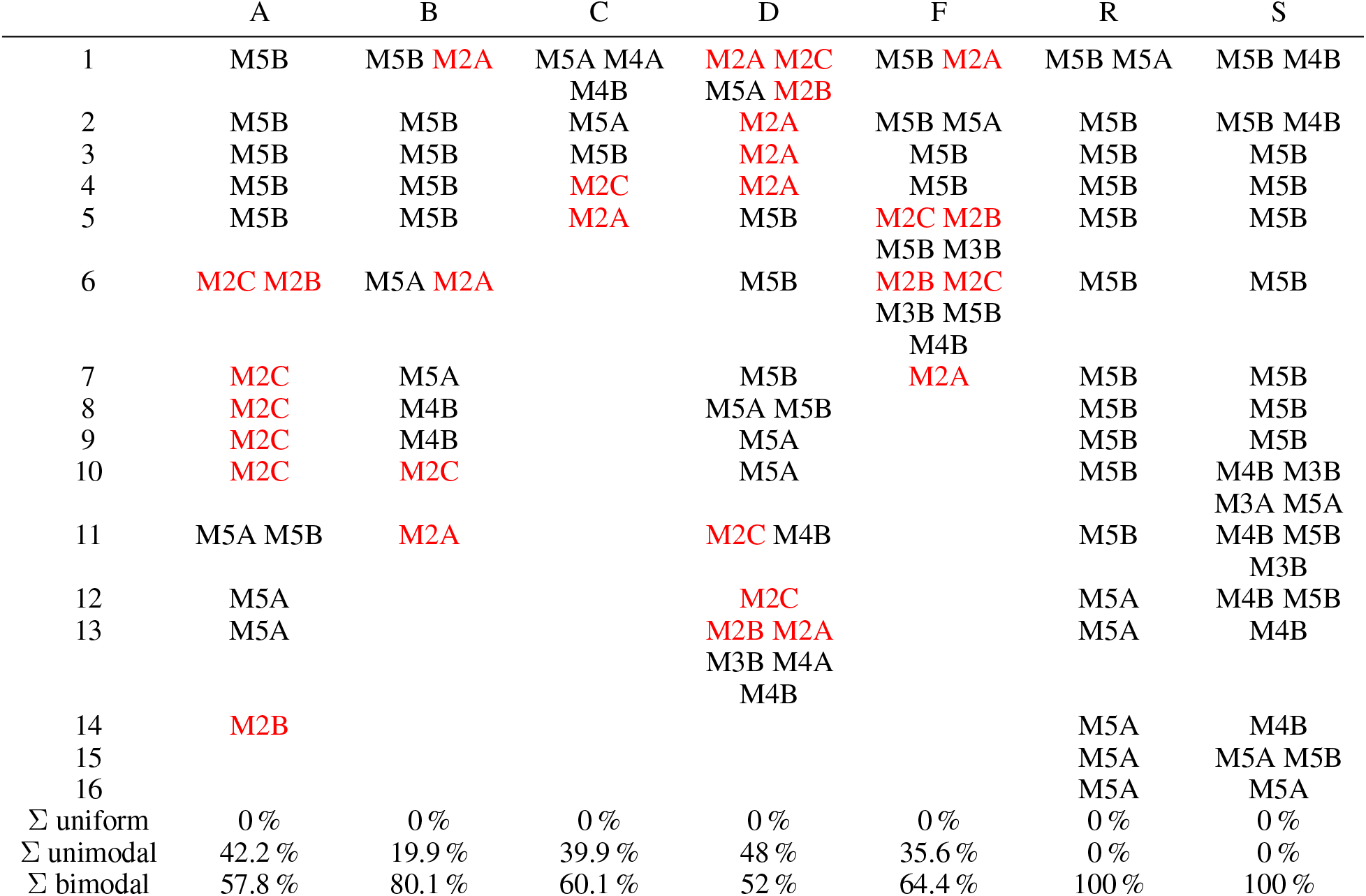
Best models for all tested bees. The table includes all test bees and 16 each of the simulated groups R and S. Unimodal distributions are depicted in red (M2A, M2B, M2C), bimodal distributions in black (M4B, M5A, M5A, M4B, M4B, M3B). Listed are all models with probability larger than 10%, ordered by decreasing probability. The last three rows give the probability of uniform, unimodal and bimodal models for each group of bees, these include those all models, even those below the cut-off of 10 %.

All distributions are non-uniform (Table 1, File S2). Bimodal distributions of directions are characteristic for the random components of the search strategy because of the design of the experiment. No unimodal distributions were found in the modeled groups, whereas all experimental groups contained different proportions of unimodal distributions (A: 7, B: 4, C: 2, D: 8, E: 4 with at least one unimodal distribution out of 16 animals). Bees from home areas from A, C and E appear to apply different search strategies, while bees from home areas B and D appear to differ most from each other. These results support the conclusion that the search strategies of the experimental groups are not fully captured by random searches although the random component appears to be an important part of their behavior. Furthermore, the groups from different home areas might differ with the strongest unimodal component for the home area group D and the lowest unimodal component for the group B indicating that the experience in the home area has an effect on the search strategies in the test area.

The frequency of directions depended to some extent on the distance between the release site and the respective fix (Fig. 4C). This is particularly relevant for the R model since the bias for reduced frequencies in the sector EES–W cannot be fully explained by the cut off of the radar in this directional sector, a feature of our experimental design taken into account by the R model. A qualitative inspection of the effect of distance indicates deviations from equal proportions, for example in test animals from home area A (e.g. distance 100m to 200 m and in those from home area B (e.g. 200–300m). These results exclude the possibility of solely random search and ask for further analyses of both the directions of stretches of search flights and the spatial distribution of fixes.

### 3.3 Quantifying the generalization and transfer effect

Next, we examined whether the spatial distributions of fixes differed between the experimental groups and the results of the two models. We constructed a 19 × 18 matrix of equally sized tiles (100 m × 100 m, numbered 0 to I in W–E and 0 to H in N–S direction, with Tile 99 centered at the release site), determined for each bee the relative number of fixes within each tile and performed pairwise comparisons for each tile between the respective frequencies. Figure 5A shows representative examples while the full data is given in File S3 (heat map per bee, as in Fig. 5A), File S4 (significance of each one of the 19 · 18 = 342 tiles, as in Fig. 5C–D), File S5 (pairwise comparison, as in Fig. 5E), and File S6 (comparison of one group to all others, as in Fig. 5F).

**Figure 5:**
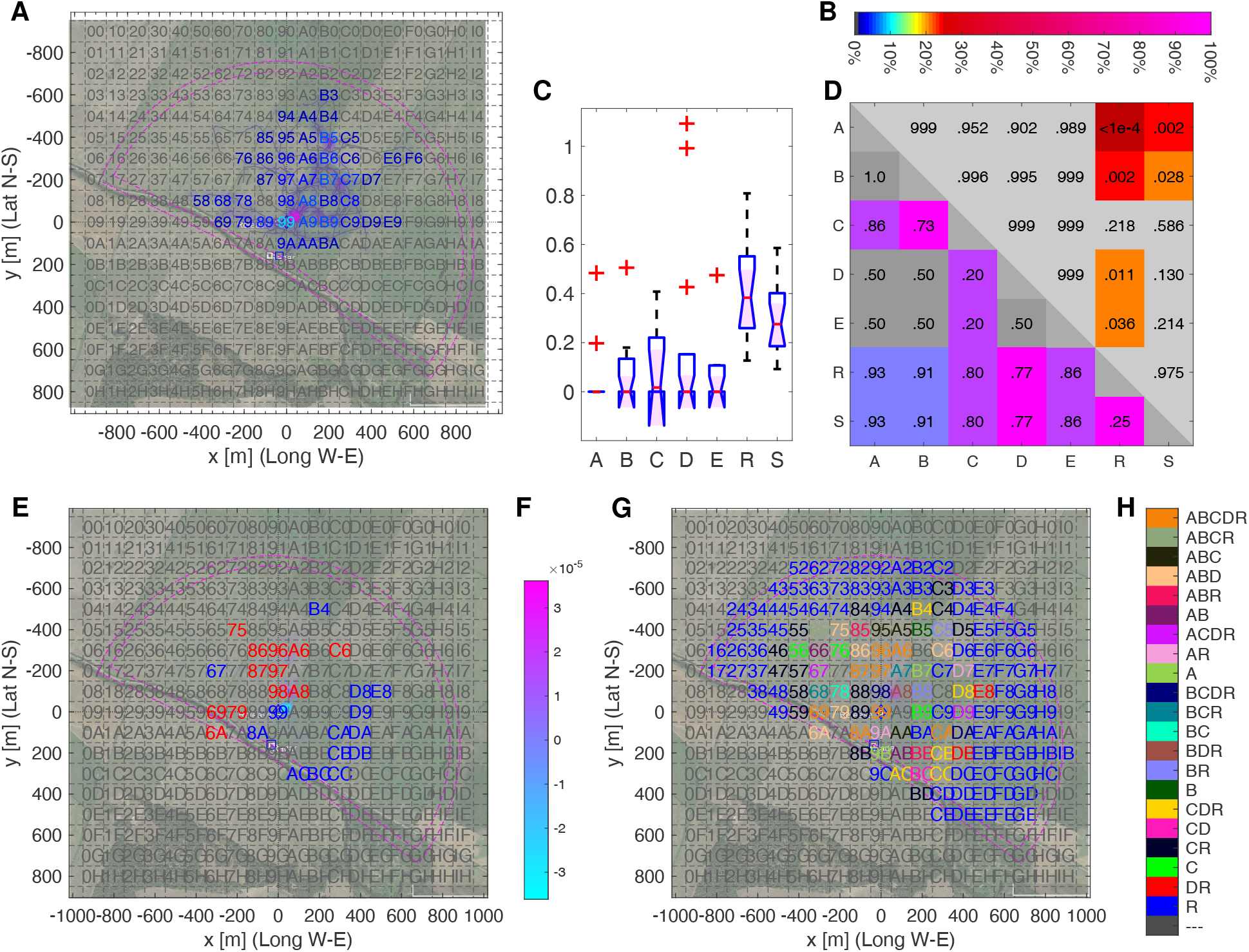
Quantification of the spatial distribution of search fixes of each individual bee from all seven bee groups. **A–B**: Frequency of fixes in the tiles of the 19 × 18 grid (each tile: side length 100 m) for Bee A5 (the 5th bee from home area A) overlaid on an aerial view of the test field. All respective frequency distributions are given in File S3. The two magenta lines surrounding the test area give the radar range (inner half circle) and the extended range 50 m further out, which corresponds to the moving average filter size (outer half circle). The heat map is generated by smoothing the fixes with a moving average filter whose size is exactly one tile. The boxes’ color represents the relative frequency in that box as coded by the colorbar in **B**, while **F** shows the heat map colorbar. **C–D**: Box 46 as one example of the pairwise comparison of each of the 342 tiles applying the Kruskal-Wallis- (D, upper half) and the MES-statistics (D, lower half), where significant values are highlighted in color. The box plots in C give the median, the inter-quartile (25% and 75%), and the outliers of the frequencies of fixes in the respective tile together with the number of test animals. The full data set is shown in File S4. **E**: Example of the pairwise comparison between bees from two different home areas (here: bees from area E compared with those from area D) using the mes value of the MES analysis (with a threshold of Δ_mes_ = 0.2). All pairwise comparisons are shown in File S5. The outer bar marks in blue frequencies of fixes of bees from area E being lower than that of bees from area D, in black frequencies lacking significant differences, and in red frequencies being significantly higher in area E area as compared to area D. The inner bar color codes the difference between the differences fixes of bees from the two areas. The edges in the test area are marked by thin pink lines. The heat map color is shown according to **F**. **G–H**: Color coded tiles (G) with color codes in H for significant differences (Δ_mes_ of at least 0.2) between bees of home area E to other bees. The outer bar color codes in the number of the respective tile the significant differences between the frequencies of fixes of bees for the given home areas. The heat map is colorized using the inner bar color codes, which depict the frequency of fixes in the area of the respective tile for bees of home area E. The frequencies of the model S are not included here. All respective calculations are given in File S6.

For statistical comparison, we applied two procedures, the Kruskal-Wallis- and the mes-statistics. Figure 5C and D shows one example of the pairwise comparison (complete data set: File S4). The result of the statistical analyses of the 342 tiles and all pairwise comparisons are composed into tables like the one shown in the example Figure 5D. A data pair deviating by Δ_mes_ ≥ 0.2 is considered significantly different. The frequency of fixes can be higher or lower in the comparison between the two groups of data. Figure 5E shows the example for bees from home area D and E. The other pairwise comparisons are given in File S5. Many of the tiles are marked with significant differences of the respective frequencies. Since 21 sheets of comparisons are hard to read, we compiled 6 summary tables of the kind shown in Fig. 5H excluding model S because this model includes a large sector that is not scanned by the radar (see File S6 for all comparisons). Here, we apply a scheme of color coded tiles that reflect significant differences between bees from one home area and all other bee groups (real and simulated). For example, in Fig. 5G, Tile B4, numbered in yellow, has as legend CDR, which indicates significant differences between the frequencies of fixes of Group E to those of Group C, D, and R, while there is no significant difference between E and those of Group A or B. Multiple significant differences were found between the distributions of frequencies of fixes both between the model calculations and the bees from the 5 home areas and between the five home areas. We conclude that the spatial distribution of fixes differs between the five home areas and cannot be explained by random flights.

To further quantify the generalization effect, we applied a variant of principle component analysis (PCA), a partial least squares regression (PLS) analysis, to our behavioral data in search for a similarity-difference gradient of the unknown guidance parameters, as shown in Fig. 6 (see also File S8 for more PLS results). PLS estimates a linear model fitting the predictor X, here the sampled heat map data (342 dimensions) for each of the 66 bees, and to the observation data Y (here the seven-dimensional, binary selector of the home area of each one of the 66 bees). In comparison to PCA, PLS’s advantages are that it can deal with correlations in the rows of the predictor matrix and that it takes Y directly into account, while PCA only works on X. Similarly to PCA, the PLS’s principal components can be used to obtain a reduced order linear model. We then analyzed the three most relevant components for each bee. We performed three different PLS analyses. The first included all seven groups (five test and both model groups R and S). Its three main components explained 91.4% of the variance. The second analysis ignored the S group, and there the three main components explained 90.6% of the variance, while the third analysis, which ignored all test bees, explained 87.8 %.

**Figure 6:**
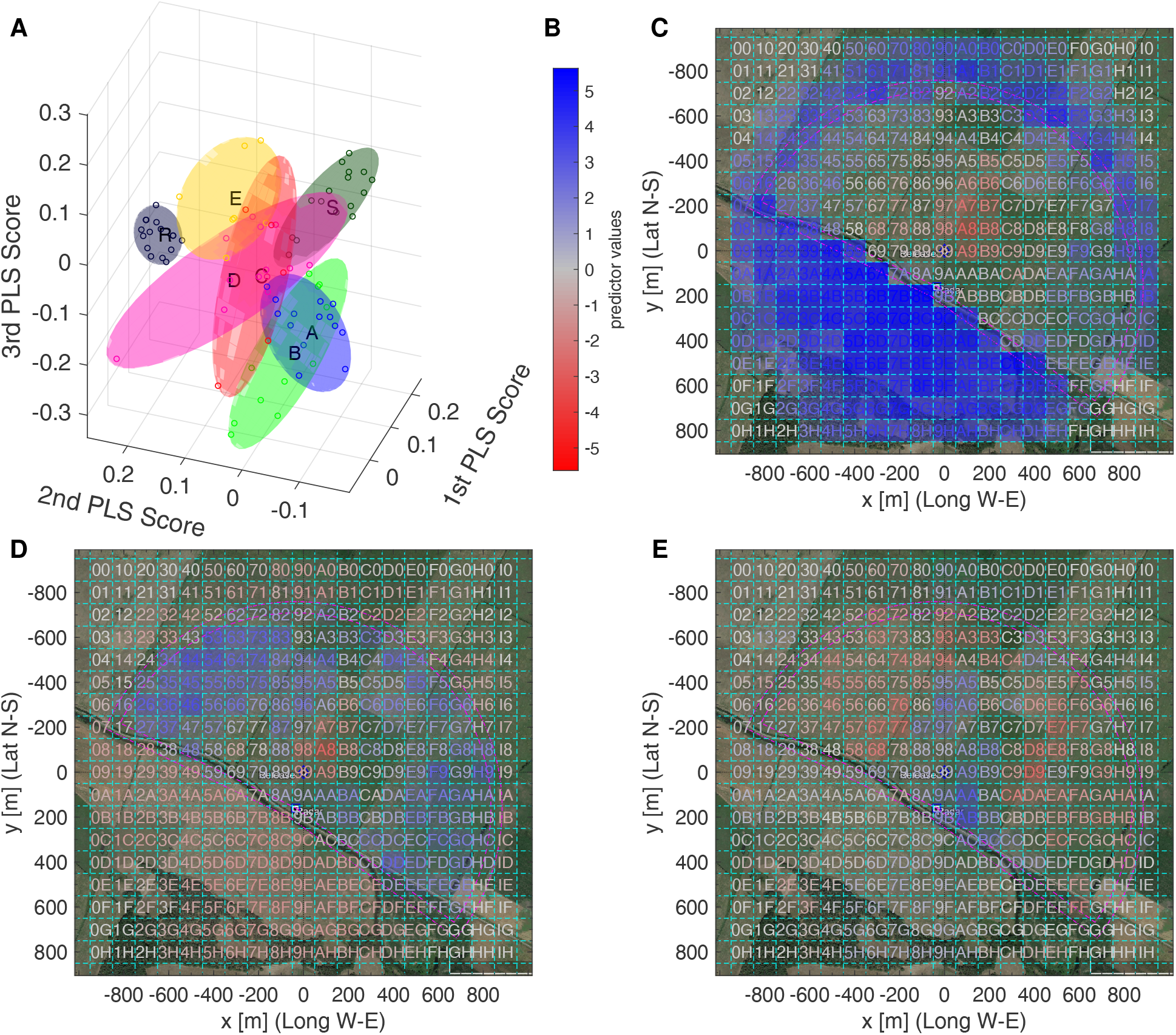
PLS analysis of the heat maps of radar fixes of all test and model groups. **A**: 3D plot of the top three PLS scores for each bee’s heat map (circles), together with minimum volume ellipsoids covering all PLS scores of one group. Color code of the ellipsoids and circles: A, B, C, D, E, R, and S. **B** Colorbar of the PLS support vectors values for C–E. **C**–**E**: Three most important PLS support vectors viewed as heat maps, with color coding as in B.**C**: First PLS support vector, **D** second, and **E** third.

The PLS scores can be visualised in 3D (Fig. 6A) and their support vectors as heat maps (Fig. 6C to E). Figure 6A additionally shows the minimum volume ellipsoids, calculated using a singular value decomposition, that contain the 3D PLS scores of each bee in a group.File S7 contains X3D-files that allow for rotating and zooming the 3D representation. As could be expected, model S is well separated (Fig. 6A) from all other groups as only its bees fly in the radar blank, but also the R model bees differ from the real bees (Fig. 6A). The latter is also clearly visible when studying the separatrix, a hyperplane in 3D that optimally separates the R model bees from all test best, in the SVM analysis that ignores the S group (Fig. 7A, E and H, see also File S9 for more separatrices). Figure 7A visualises this separatrix as heat map, showing that the R bees fly more frequently in the red heat map boxes than the other bees. One of these red boxes is Box 46, whose statistics is shown in Fig. 5C to D. The signed distance of each bee to the heat map (Fig. 7E), where positive and negative value correspond to the SVM classification into two classes, shows that all R bees are above (positive distances) and all other bees below (negative distances) this separatrix, with the separatrix itself shown as dashed line. The corresponding statistics shows very significant differences between R bees and each other group, but also between any two of the test bee groups (Fig. 7H).

**Figure 7:**
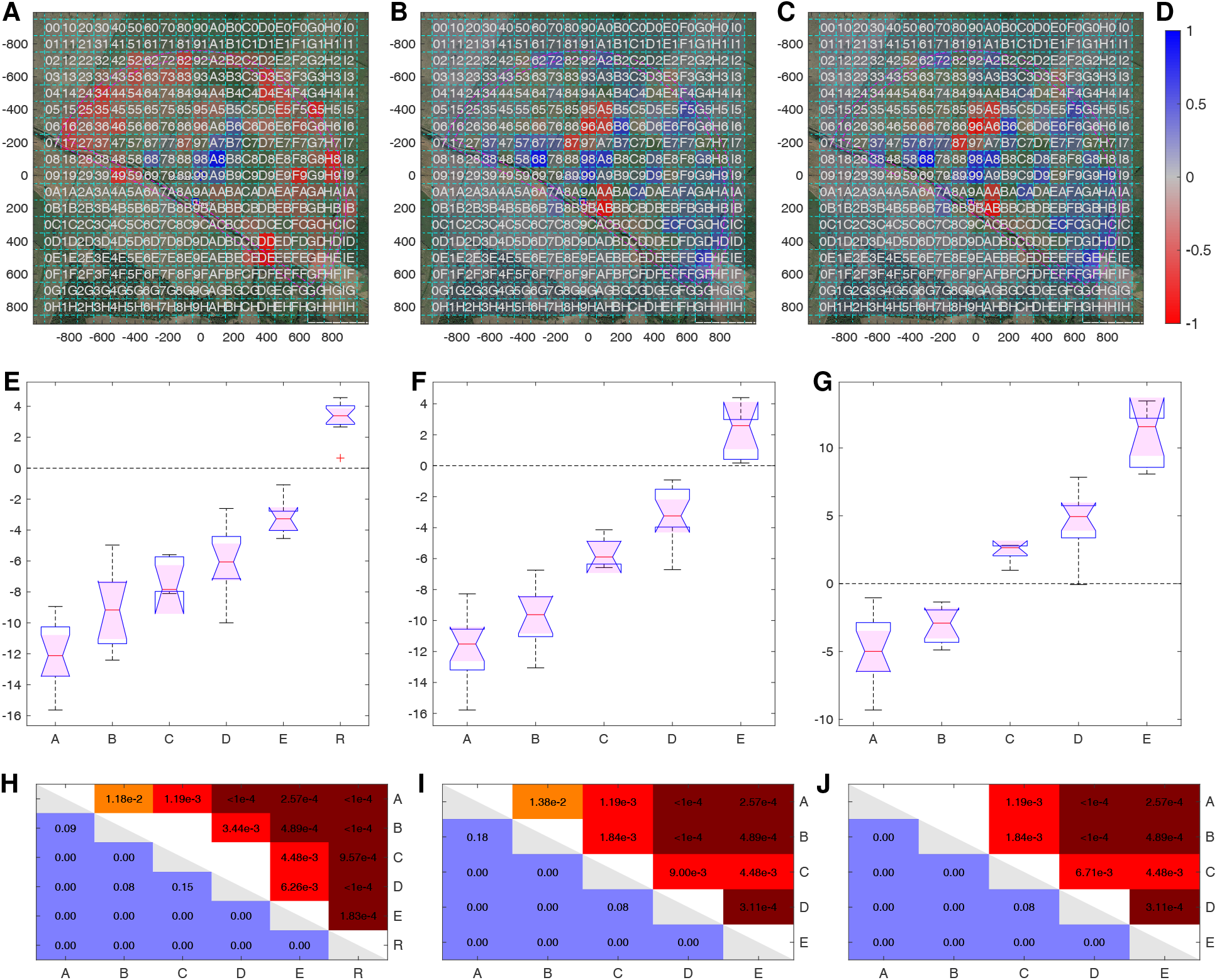
Separatrices, obtained using SVM, of the three main PLS components. **A**, **E**, **H**: hyperplane separating R bees from all other groups, with PLS that ignored S group. **B**, **F**, **I**: hyperplane separating C and D groups from the E group, with PLS that ignored R and S group. **C**, **G**, **J**: separating A and B groups from the C and D groups, with PLS that ignored R and S group. **A–C**: Separatrices viewed as heat maps. **D**: Color code for the boxes in A–C. **E–G**: Boxplot of the distance of each bee to the separatrix, sorted by group. **H–J**: Statistics (mes lower left, Kruskal-Wallis upper right).

The SVM analysis ignoring the test bees shows even more clearly that the test bees can be grouped into three categories: A and B, C and D as well as E. Figure 7 exemplarily shows two other separatices. The middle column shows the separatrix that separates Group E from Groups C and D, while the A and B group are not considered when calculating this separatrix. Group E is fully above and all other bees below the separatrix (Fig. 7F). Also here, there are significant statistical differences between any two pair of bee groups (Fig. 7I). The corresponding heat map reveals that E bees searched more to the north and south of the Release, and less in several other spots as e.g. in the south-east, colored in blue (Fig. 7B).

The third separatrix shown in Fig. 7,C, G and J was defined to optimally separate Groups A and B from Groups C and D. Also here, a perfect separation was obtained (Fig. 7G), with Group E finding itself completely on the side of the C and D groups, with very significant pairwise differences (Fig. 7J), except between Groups A and B when using Kruskal-Wallis. The heat map (Fig. 7C) is very similar to the previous one(Fig. 7C), which shows that the angle between these two separatices is relatively small. Notably, the PLS analysis depicted in Fig. 7 shows that bees from home area A and B fly differently to those of C and D and all the four also differently from those of home area E, with a dominant trend from A to E and R in alphabetical ordering.

The previous analyses completely ignored the landscape of the test area. The next section studies how the bee flight paths correlate to the dominant landscape structures of the test area.

### 3.4 The guiding effect of edges

The common test area lacked a skyline of an angular modulation of > 2° visual angle for the NW, N, NE and E directions and contained a row of bushes in the SE, S and SW directions that presented angular modulations of > 2° visual angle when the test bees came closer to these bushes than the release site. Interestingly, homes areas A, B and C had rather similar patterns of edges as that in the test area but tilted with respect to N. D and F did not contain extended edges, and F had no edges as common in agricultural areas but two crossing forest aisles. Thus, the test area differed from all 5 home areas with respect to the skyline pattern and the distribution of rising local landmarks. In contrast, the test area’s elongated ground structures, called edges here, of the test area (two narrow creeks NW and NE of the release site, and an edge of differently cut grass running approximately S–N close to the release site) may well partly resemble edges in the home area, particularly to those in home area A and B, slightly in C, but not at all in D and E (Fig. 1). Deviation from random search and differences between the 5 groups, thus probably reflects a generalization effect due to different layouts of their respective home area, as all bees were subject to identical experimental procedures. We first focused on the relative time each bee spent near one of the four edges, and then on the bees’ direction of flight relative to the edge direction when close to these edges. For each bee, the radar pixels were interpolated, see Section Methods, and for each resampled flight pixel, the closed edge in a Euclidean distance sense was determined. Exemplarily, Fig. 8A and C show two bees’ flight paths with color highlighting of points close to one of the four edges, with the color intensity coding the distance range (Fig. 8B). All bee paths are shown in File S10. From Fig. 8A and particularly Fig. 8C it is clearly visible that many fixes are further away than 100 m to the closest edge (path as thin gray line). The relative time near an edge was then quantified as relative number of pixels that are at most 10 m, 25 m, 50 m, or 100 m from their closest edge. Figure 8D and E show this for the Bees A05 and R01, respectively, for the others see File S10.

**Figure 8:**
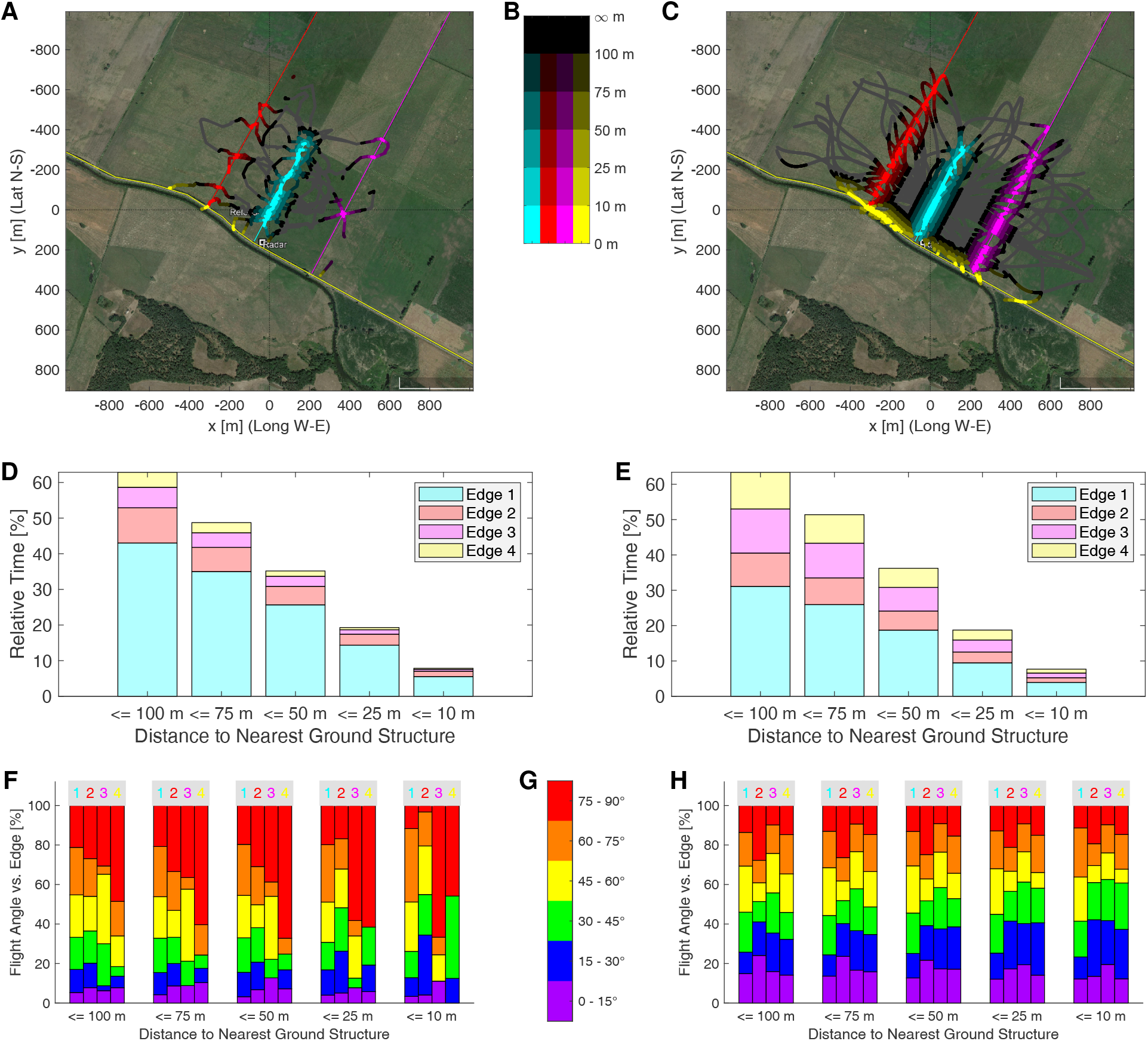
Near edge analysis. **A**–**C**: Flight path of Bee A05 (**A**) or Bee R01 (**C**) with **B** showing the color-code used for the distance to nearest edge up to at most 100m. The full path is plotted as a thin gray line. **D**–**E**: Relative time spent near an edge for Bee A05 (**D**) and Bee R01 (**E**), evaluated in five distance ranges (up to 100m, 75m, 49m, 25m, or 10m). **F**–**G**: Relative time spent in an angle sector relative to the nearest edge, Bee A05 (**F**) and Bee R01 (**G**), evaluated in the five distance ranges of **D**–**E** and six angle ranges of 15° width. Each bar corresponds to one edge and one distance range, the color bars’ height relates to the relative time spend in the corresponding angle sector.

As bees have been shown to “see” edges from approximately 30m (Menzel, Tison, et al., 2019), we restrict in Fig. 9 the analysis to the distance range of at most 25 m. The distribution of the relative time spent per bee at this distance range shows a large heterogeneity among the test bees, while the model bees are much more homogeneous. Applying the Kruskal-Wallis tests only, few significant differences are found and only between model bees R and the test bees A, B, and D, when using Edge 4, and only to bees A for Edge 1 and 3. However, Cohen’s U3 mes highlights significant differences in many more cases. Bees A differ from all others with respect to the time spent near Edge 1, the edge passing close to the release site. Bees A also spent more time there. Edge 3 is the opposite, bees from home area A spent significantly less time there than bees from other hives and also less than the model bees R. The latter also differ from Bees C and E as the model bees spent more time close to Edge 3. The heterogeneity among Bees B prevents the test from providing any significant difference between Bees B and any other group. Edge 2 is useful for discriminating Bees E, as these bees significantly less time there as bees from hive C and D as well as the model bees R. Interestingly, there is a high heterogeneity among bees A, B, and C with respect to the amount of time spent near Edge 2. Edge 4, the bushes behind the radar, give quite some discrimination information, even though the bees flew relatively rarely there. The mes test shows significant differences between 11 out of 15 pairs. Only the pairs of bees A–B, C–E, C–R, and E–R show similar closeness. Overall, using all four edges allows for discriminating any pair of bee group.

**Figure 9:**
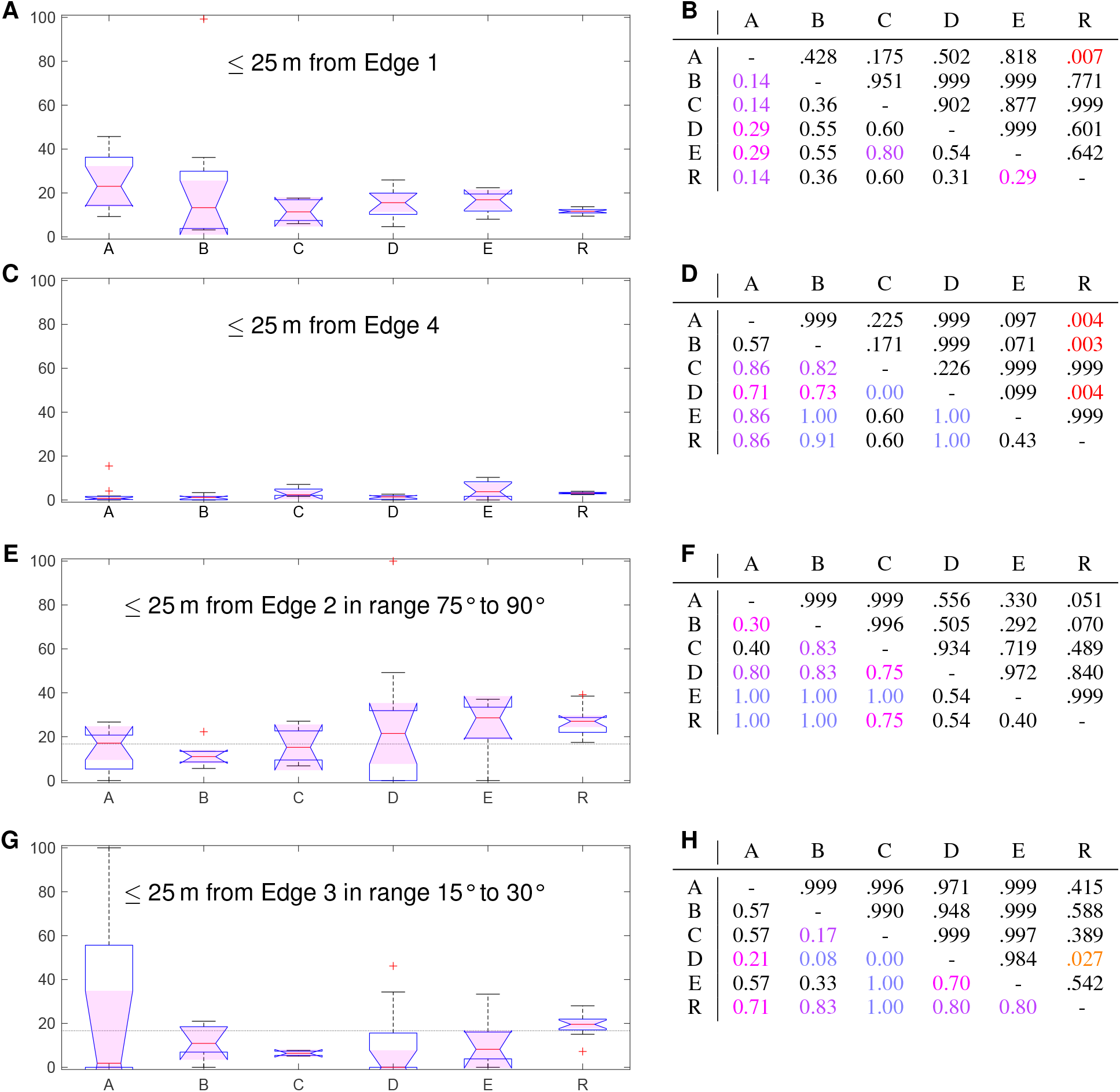
Relative time spent nearer than 25 m from edge. **A–B**: Near Edge 1. **C–D**: Near Edge 4. **E–F**: Near Edge 2 with flight angle relative to edge in 75° to 90°. **G–H**: Near Edge 3 with flight angle relative to edge in 15° to 30°.**A, C, E, G**: Boxplot of values per bee, sorted by bee group, **B, D, F, H** Kruskal-Wallis and mes-statistics.

Next, we studied the flight angle relative to the nearest edge. For each distance range as above, we counted the angles in six ranges of 15° width, i.e. 0° to 15° up to 75° to 90°. The 120 statistics, four edges à five distance ranges à six angle ranges, can be found in File S11. Figure 9C and D show two examples. The R model bees are quite homogeneous and have values usually close to the uniform distribution (dotted line at 100 %/6 ≈ 16.7 %), but with a significantly higher variance than in the time near edge, which is due to the stochastic nature of their flight paths, as the tangent of the model bees’ local turning rate is drawn from normally distributed random numbers. The test bees are in some cases very homogeneous, e.g. Group B in Fig. 9E or Group C in Fig. 9G. In most cases however, there is a significant variability within the bees of a group, e.g. Group D in Fig. 9E or Group A in Fig. 9G. The examples in Fig. 9 show that for each pair of group there exists a flight measure that is able to discriminate that pair with high mes significance.

Figure 10 summarizes the mes-statistics for the pairwise discrimination using all 20 time near edges and the 120 flight angle near edges. The Δmes of all 20 time near edges for each pair of bee groups (Fig. 10A) shows that Edge 4 is a good discriminant for many pairs, i.e. B–C, C–D, D–E, D–R, while Edge 3 discriminates well E–R and Edge 1 the pair A–B. Having a closer look at the best and second-best time near edge measurements shows that the time near edges alone allows in almost all cases for very good discrimination (of at least Δmes = 0.43, often even equal to the maximally attainable Δmes = 0.5) (Fig. 10C). Only the pairs of bees A–D, B–D and C–E have lower values, with B–D the lowest at 0.23. The time spent near Edge 2 and 3 showed much less significant differences than near Edge 1 and 4 (File S11).

**Figure 10:**
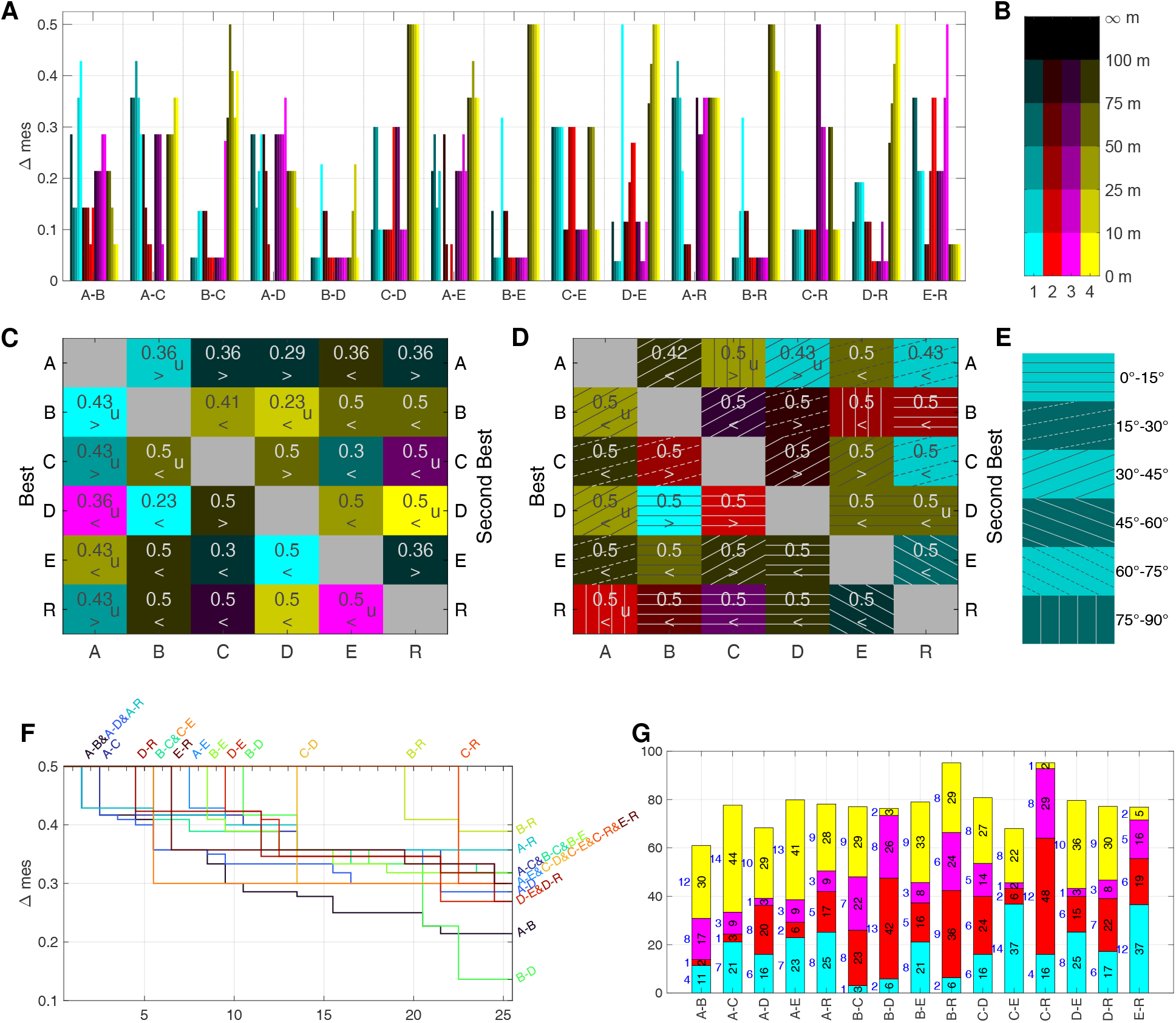
Discrimination analysis using the mes statistics based on time and flight angle near edges. **A**: Δmes of time near edge of each pair of bee groups, for each of the four edges (color) and five distance ranges (color intensity). Color coding of closest edge and distance to edge as in B. **B**: Legend for the distance range to closest edge. **C**: Best discriminating cases from A (lower left) and second best (top right); the greater/smaller sign means the row label has larger/smaller values that the column label. A “u” denotes those best (respectively second best) cases that a unique, i.e. having a larger Δmes than the second (respectively third) best case. **D**: Same as C, but for the flight angles, the angle ranges are coded by the stripe pattern shown in E and colorised as in B. **E**: Legend for the flight angle ranges, coded as stripes. **F**–**G**:Δmes for the 25 best discrimination measures for each pair of bee groups, combining time near edges as well as angles.**F**:Δmes values (y-axis), sorted in decreasing order, with index 1 to 25 on the x-axis.**G**: Number (in blue on the left of each bar) and sum (black and height of bar) of best Δmes values per group pair and edge. The height is scaled to 100 %, which corresponds to 12.5, i.e. 25 times the maximal Δmes of 0.5.

Often, the Δmes of a pair of group at various distances to one of the edge are similar or even equal, as e.g. for Edge 4 in C–D, B–E ((Fig. 10A). This is, however, not always the case as Edge 1 in B–E and D–E shows where the fixes close than 10 m have a discrimination of Δmes = 0.5 while all ranges further away from Edge 1 do not show a significant discrimination. In some cases, the ranges close to an edge show higher discrimination value, e.g. Edge 3 in E–R or the already mentioned case Edge 1 in B–E and D–E. In others, the ranges further away from the Edge are more discriminatory than the closer ones, see for example Edge 4 in C–E, Edge 3 in C–R, Edge 1 ind E–R.

The flight angles allow for an even better discrimination, as each pair has at least one Δmes = 0.5, most pairs have several test cases maximally discriminating between them (Fig. 10D). Even better discrimination arises by using a combination of time near edge and angle near edge test cases. Figure 10F shows for each pair of bee groups the values of the best Δmes of time and angle near edge test cases. For any of these pairs, more than 20 test cases exist having a Δmes > 0.2, thus showing a significant difference. Also, all pairs can be discriminated with at least five test cases having a Δmes > 0.3. Two pairs are particularly distinctive: both B and C bees 19 or more test cases that are maximally distinctive. This shows that their search strategies are very different from random searches. While the other test bees can also be well discriminated from the R group, the number of test cases highly discriminating them from the R group is much lower. This is a notable result as the pairs A and B as well C and D seemed to behave similarly in the heat map analysis.

An interesting result of the edge analysis in Figure 10C and D uncovered that the statistically relevant zone is not only close to the edges, but also beyond the distance bees were shown to see them, i.e. approximately 30 m (Menzel, Tison,et al., 2019).

Figure 10G highlights which edges are present in the top 25 discriminating test cases shown in Fig. 10D. The bar heights are proportional to the cumulated Δmes per pair while the colored bar with the number to their left show how many of the 25 test cases are near one of the four edges, with Edge 1 in cyan at the bottom and Edge 4 in yellow at the top. All Edges are present for each pair, however differences are clearly visible. For example, Edge 1 is not important for discriminating the pairs B–C and B–D, but plays a significant role for discriminating E from C, D, and R as well as A from C, E and R. Edge 2 and 3 are extremely important for the pairs B–D and C–R, and also much for B–C and E–R. Edge 4 plays a significant role for most pairs, with notable exception of the pairs B–D, C–R and E–R.

We conclude from the analyses above that bees from the different home areas were guided in the test area by a generalization effect of their navigational memory. The parameters of guidance are not obvious, although it appears that the existence or lack of edges in the home area (their visual features, their extension in the compass direction, their straightness, and other unknown features) might play an important role. An additional discriminating parameter could be the lack of a panorama in the test area. The aerial views of the homes areas (Fig. 1) shows more or less similarities with these landscape features with those in the test area. Home areas A and B were more similar than C to F, and home areas E and F were most different from the test area. We hypothesize that the edges in the test area provide guiding structures that may be differentially used by bees from the different home areas. For example, the A bees fly more around Edge 1 than the other bees, while A and B bees stay away from Edge 4 compared with the other bees. Comparing home areas A and B, one would expect that Bees B fly less near Edge 2. This is indeed the case for the majority of them, a part of these bees however spend a significant amount of time near Edge 2. This could be explained by Edge 2 possibly resembling the IRC close to their hive in home area B.

## 4 Discussion

Successful navigation requires forming a lasting memory of the locations and identities of significant objects in the environment in relation to a compass. This memory needs to be updated continuously when the animal moves allowing effective and flexible navigation. Multiple perceptual systems are involved in probing the world during navigation, and vision is usually the most important sense in further reaching navigation. Recognizing and storing the spatial relations of objects requires reference systems of two kinds, egocentric and environmental (or allocentric). Egocentric references include view point memories, path integration (or dead-reckoning) and body relations to a geocentric reference like the sun compass (T. S. Collett and Rees, 1997;T. S. Collett, Graham, and Durier, 2003; Wehner, Michel, and Antonsen,1996). Environmental (allocentric) references structure memory such that the spatial relations between egocentric and geocentric references as well as the spatial relations between identified objects are stored. The level of integration between egocentric and allocentric references in insects, and particularly in honeybees, is under debate (T. S. Collett and Graham, 2004). The underlying neural processes may be conceptualized as the activation of multiple isolated ad hoc procedures or as the retrieval of a concise navigation memory. In the latter case generalization and transfer phenomena may provide us with hints about the level of integration across the multiple neural processes involved. Support for this view comes from multiple observations in test conditions in which close visual cues at a feeding site were systematically changed both during training and testing in order to uncover higher order memory processing (Giurfa, 2003; 2015). In these experiments, bees were asked whether they generalize across learned cues that can be discriminated but contain hidden parameters that binds them to categories, i.e. learning of bilateral symmetry (Giurfa, Eichmann, and Menzel, 1995), matching-to-sample (Giurfa, Zhang, et al., 2001).

We have used here a generalization test procedure to explore the properties of landscape memory. In learning theory generalization is the counter part of discrimination, the usual test procedure in navigation experiments. In discrimination experiments animals are trained to distinguish between differentially trained stimuli. In generalization animals are asked whether and how they transfer what they have learned to other stimuli they are able to discriminate. The transfer is either immediate and specific, or used later for new learning.

We deal here with immediate specific transfer, the ability of an animal to respond to partially learned and novel stimuli in the navigation context. Based on the rich literature about navigation in bees (see Introduction) we believe that the existence or absence of panorama and local cues, the kind of panorama and local cues as well as the existence, appearance and compass directions of edges are very well learned and discriminated by bees. However, this was not verified for the cues involved in the experiments reported here. In particular the learning and discrimination of isolated cues involved was not studied. Experiments under natural conditions and within the dimensions of natural navigation training procedures are close to impossible because environmental features cannot be moved around systematically, separated or freely combined. This is unfortunate because learning about a stimulus or combinations of stimuli influences strongly how the behavior is generalized to other stimuli. In laboratory experiments different training procedures are used to distinguish between the animal’s ability to discriminate, and subsequently generalize to other stimuli (Blough, 1975; Kehoe, 2008). Still, generalization tests are highly informative because it is reasonable to assume that self-training and exploratory learning under natural conditions will be rather similar and close to optimal across different animals of the same colony that are exposed to the same environment. Generalization is embedded in the life history of the specific species. In our context, bees gain from generalization when environmental conditions alter the landscape features or when colonies move to a new nest site during swarming. Furthermore, the ability to generalize (rather than to discriminate/not discriminate) is thus a key component of cognition bound to phenomena like selective attention, expectation, categorization, similarity judgments and saving (Blaisdell, 2008; Rescorla, 1992; Zentall et al., 2008). “Generalization occurs in learning, and is essential for deriving knowledge from experiences and for skills of all kinds. It is the basis of predicting future situations from past experience and for drawing analogies.” (Gregory and Zangwill, 1987, p. 284). Thus, generalization speaks to the cognitive dimensions of using memory for solving a problem.

Here, foraging bees were collected from hives located in different country sites, their explored home area. They were trained to a close feeding site, collected when leaving the feeder and transported to a common test site where they were released at the same release site. The home areas were selected such that they differed more or less from the test area. In particular, the home areas provided a more or less rich panorama, and the test area lacked a prominent panorama. The test area provided mostly elongated ground structures (called edges here) of rather simple geometry, and the home areas differed in a graded way in this respect. The search flights of the test bees were recorded by harmonic radar over a scanning range of 900 m radius ensuring that natural dimensions of honeybee navigation were tested. If the test bees would not, at least partially, generalize their site specific navigation memory, they are expected to apply a stereotypical search strategy and no differences would appear between bees from different home areas. Unexperienced bees or bees unfamiliar with a landscape perform multiple exploratory loops in different directions and over different distances (Capaldi et al., 2000; Degen, Hovestadt, et al., 2018; Degen, Kirbach, et al., 2015; Menzel, Greggers, et al., 2005). Experienced bees released in an unexplored area also perform rather regular loops of search flights that have a number of common features. Bees fly in different directions over different distances during the outbound flight components and return to the release site multiple times (Menzel and Greggers, 2015). Thus, the lack of memorized guiding cues are thus likely to induce random search possibly similar to those of the well-studied desert ant *Cataglyphis* (Wehner and Srinivasan, 1981). In addition, stereotypical flights according to innate responses to landscape features may be apparent. In both cases no differences between bees of the different home areas are expected. Two models were run, one that included all directions around the release site (Model S) and one that excluded flights in the not scanned sector of the radar (Model R). Although random components certainly contribute to the search strategy of the test bees random search as the only or dominant strategy can be rejected on the base of several findings.

1. The **distribution of flight directions** differs significantly between test bees and the random models as well as between bees from different home areas (Figs. 3 and 4). This is supported by non-parametric likelihood-based models analyzing the directions of flight stretches, which uncover significant differences between all 7 groups, the 5 groups of different home areas and the two model groups (Fig. 4 and Table 1).
2. The **spatial distribution** of flight fixes quantified by heat maps reveal significant differences between any two groups of test and model bees at specific heat map quadrants (Fig. 5). A unified view was obtained by a partial least squares regression (PLS) analysis that uncovered structural differences in the flight fixes’ spatial distribution. The three dominant PLS support vectors explain approx. 90% of the heat map data variance. As PLS support vectors as well as hyperplanes separating the three dominant PLS support vectors of bee groups can be plotted as heat maps, the PLS analysis is also a highly informative way of visualizing these differences Figs. 6 and 7). Figure 7A–C show three examples of separating hyperplanes. These hyperplanes separate bee groups very well, especially the model bees R and S from all others as well as from each other, but also all test bee groups from each other with very high significance (Fig. 7H to J). Most importantly, a gradient of similarity can be derived from the hyperplane separating Group R from all experimental groups (Fig. 7A, E,H). Other hyperplanes are able to separate the test bees into three supergroups: Groups A and B, C and D, as well as E (Fig. 7E to G).
3. The previous analyses did not take into consideration the **edges** of the test area. From a view point of existence or absence of edges and structured panorama, D and E home areas differ more strongly than A and B homes areas from the test area, and home area C lies in between (Fig. 1). A and B home areas are characterized by elongated ground structures but in different ways (irrigation channel, rows of trees, compass direction). These differences are reflected in the preference of Edge 1 by A bees, less guidance by Edge 2 in B bees, and no attraction to Edge 4 for Groups A, B, and D. Their respective home areas were characterized by further distant panorama, border lines of agricultural fields and segments of elongated ground structures that differed quite considerably from those in the test area. The edges in the test area impact the search flights of the bees in each group differently (Figs. 8 to 10). Quantifying the bee’s proximity to the ground structures revealed that the bees of each groups differed significantly from each other in their flying behavior, both with respect to the time spent close to each edge (Fig. 10A and C), as well as to the angle they flew to or away from these edges (Fig. 10D). This effect was not only visible in close proximity to an edge, but also in the range of 75 m to 100 m, where the bees probably cannot see the edge. Groups B and C differ most from the random bees, see Fig. 10F. Groups B differed mostly around Edge 2 to 4 from the model bees, for Group C only near Edges 2 and 3 as well as 1, but not Edge 4. A summary of the discrimination of bee groups using the edges is not as simple as for the heat maps. Nevertheless, it is obvious from Fig. 10 that restricting the bee data to the proximity of the edges is sufficient to highly discriminate among any two pair of bee groups. This requires all four edges as for any edge there are pairs best discriminated using that edge, and other pairs, where a specific edge does not show discriminatory information.

Taken together, sole guidance by a random search strategy and the effect of stereotypical potentially innate guiding factors can be rejected. The heat map PLS analysis (Figs. 6 and 7) supports the conclusion that a similarity gradient based on the elongated ground structures guided the search flights. Bees from home area E that lacked any similarity with the test area behaved most closely to the modeled R random bees. Bees from home areas A and B behaved most different from modeled R random bees, and were close to each other. Bees from home area C and D were in between and were also rather close to each other. A detailed analysis of the effect of the edges on the different bee groups uncovered that the edges impacted the bees, both with respect to the time spent near an edge, as well as to their flight directions.

The latter finding is important because elongated ground structures might be attractive innately to bees, and flight directions might be bound to directions of elongated ground structures. Indeed elongated ground structures are frequently used for guidance in honeybees (Menzel, Tison, et al., 2019) and bumble bees (Brebner et al., 2021). These structures need to be learned (discussion in Menzel, Tison, et al., 2019). Elongated ground structures like irrigation channels, rows of bushes or trees, edges of forest and a river bank characterized the home areas differently. Two irrigation channels and a borderline between two grasslands were the only elongated ground structure in the test area. A row of bushes and a parallel running small road as well as a parallel creek were in the south of the radar. Most of the flights beyond the row of bushes was, however, occluded by the radar blanking sector. The skyline of the horizon was even over most of the test area covered by the searching bees within the radar range (≤ 2° visual angle). This was not the case in any of the 5 home areas. Thus, the effect of panorama matching was not addressed in our experiments, but the lack of any panorama cue in the test area may have impacted the search flights of the test bees differently. Piloting towards a beacon can also be excluded because the only beacon in the test area was the radar antenna that was not approached. Preferred compass directions can also be excluded for both the earth magnetic field and the sun compass. In both cases such preferred directions would have to be detected in all 5 experimental groups. This applies also for the possibility that searching bees would prefer a constant angle to the sun azimuth because the experiments were performed between a constant time window (12:00 am and 3 pm). We thus conclude that the home area specific effects indicate a generalization effect of navigation memory acquired in the respective home area.

Elongated ground structures are characterized by unique features for flying animals that make them most suitable as reference objects in mid-range navigation. They keep a stable relationship to a compass direction (Dyer and Gould, 1983; Frisch and Lindauer, 1954), they are polarized in relation to the goal (leading to, leading away) and in relation to other localized landmarks, they may be identical to elongated landmarks at the goal, thus following it will lead the animal to the goal, they provide potentially a network of extended landmark features that characterize locations uniquely. These alignment effects has been well studied in human navigation (McNamara, Rump, and Werner, 2003). In laboratory mammals, the alignment effect requires a functional hippocampus, possibly via boundary vector cells (Barry et al., 2006). Robots were found to use line-like landmarks for efficient navigation (Furlan, Baldwin, and Klippel, 2007; Se, Lowe, and Little, 2005). Most importantly, flying animals identify such extended ground structures in a map-like aerial view making them highly attractive as guiding structures. It is thus not surprising that both bats and birds use linear landmarks for navigation (Biro, Meade, and Guilford, 2004; Geva-Sagiv et al., 2015; Heithaus, Fleming, and Opler, 1975; Lipp et al., 2004). Based on the data reported here we conclude that elongated ground structures are also salient components of the honeybees’ navigation memory.

## Acknowledgements

The authors appreciate Monica Schliemann-Bullinger’s valuable and profound comments to various drafts of this manuscript.

## Author Contribution

**Conceptionalization** Uwe Greggers, Randolf Menzel

**Statistical Analyses** Eric Bullinger

**Writing** Eric Bullinger, Randolf Menzel

## Supplementary Material

The files listed below can be found on the OSF plattform.

**File S0** Primary data

**File S1** Directional analysis

**File S2** Circle statistics

**File S3** Heat maps

**File S4** Significance analyses of heat map tiles

**File S5** Pairwise comparison of bee groups wrt. heat maps

**File S6** Significant tiles for comparing one bee group to all others

**File S7** PLS result in 3D

**File S8** PLS results as heat maps

**File S9** PLS separatices as heat maps

**File S10** Flight paths near edges

**File S11** Significance analyses of time and flight angle near edges for all bee groups

## References

Akaike, H. (1992). “Information theory and an extension of the maximum likelihood principle. Foundations and basic theory”. In: Breakthroughs in Statistics. Ed. by S. Kotz and N. L. Johnson. Vol. 1. New York: Springer, pp. 610–624.

Akaike, H. (1973). “Information theory and an extension of the maximum likelihood principle”. In: Proceedings of the 2nd International Symposium on Information. Ed. by B. N. Petrov and F. Csaki. Reproduced in Akaike, 1992, pp. 267–281.

Barry, C. et al. (2006). “The boundary vector cell model of place cell firing and spatial memory”. In: Reviews in the Neurosciences 17.1-2, pp. 71–97.

Batschelet, E. (1981). Circular Statistics in Biology. London: Academic Press.

Biro, D., J. Meade, and T. Guilford (2004). “Familiar route loyalty implies visual pilotage in the homing pigeon”. In: Proceedings of the National Academy of Sciences 101.50, pp. 17440–17443.

Blaisdell, A. P. (2008). “Cognitive dimension of operant learning”. In: Learning and Memory: A Comprehensive Reference. Ed. by J. H. Byrne and R. Menzel. Vol. 1. Oxford: Academic Press. Chap. 10, pp. 173–195.

Blough, D. S. (1975). “Steady state data and a quantitative model of operant generalization and discrimination”. In: Journal of Experimental Psychology: Animal Behavior Processes 1.1, pp. 3–21.

Brebner, J. S. et al. (2021). “Bumble bees strategically use ground level linear features in navigation”. In: Animal Behaviour 179, pp. 147–160.

Capaldi, E. A. et al. (2000). “Ontogeny of orientation flight in the honeybee revealed by harmonic radar”. In: Nature 403, pp. 537–540.

Cartwright, B. A. and T. S. Collett (1983). “How honey bees use landmarks to guide their return to a food source”. In: Nature 295, pp. 560–564.

Cheeseman, J. F. et al. (2012). “General anesthesia alters time perception by phase shifting the circadian clock”. In: Proceedings of the National Academy of Sciences 109.18, pp. 7061–7066.

Cohen, J. (1988). Statistical Power Analysis for the Behavioral Sciences. Cambridge: Academic Press.

Collett, M., D. Harland, and T. S. Collett (2002). “The use of landmarks and panoramic context in the performance of local vectors by navigating honeybees”. In: Journal of Experimental Biology 205.6, pp. 807–814.

Collett, T. S. and J. A. Rees (1997). “View-based navigation in hymenoptera multiple strategies of landmark guidance in the approach to a feeder”. In: Journal of Comparative Physiology A 181, pp. 47–58.

Collett, T. S. and P. Graham (2004). “Animal navigation: path integration, visual landmarks and cognitive maps”. In: Current Biology 14.12, R475–R477.

Collett, T. S., P. Graham, and V. Durier (2003). “Route learning by insects”. In: Current Opinion in Neurobiology 13.6, pp. 718–725.

Collett, T. S., N. H. de Ibarra, O. Riabinina, and A. Philippides (2013). “Coordinating compass-based and nest-based flight directions during bumblebee learning and return flights”. In: Journal of Experimental Biology 216.Pt 6, pp. 1105–1113.

Degen, J., T. Hovestadt, M. Storms, and R. Menzel (2018). “Exploratory behavior of re-orienting foragers differs from other flight patterns of honeybees”. In: PloS One 13 (8), e0202171.

Degen, J., A. Kirbach, et al. (2015). “Exploratory behaviour of honeybees during orientation flights”. In: Animal Behaviour 102, pp. 45–57.

Dyer, F. C. (1996). “Spatial memory and navigation by honeybees on the scale of the foraging range”. In: Journal of Experimental Biology 199.Pt 1, pp. 147–154.

Dyer, F. C., N. A. Berry, and A. S. Richard (1993). “Honey bee spatial memory use of route-based memories after displacement”. In: Animal Behaviour 45.5, pp. 1028–1030.

Dyer, F. C. and J. L. Gould (1983). “Honey bee navigation”. In: American Scientist 71.6, pp. 587–597.

Fitak, R. R. and S. Johnsen (2017). “Bringing the analysis of animal orientation data full circle: model-based approaches with maximum likelihood”. In: Journal of Experimental Biology 220.21, pp. 3878–3882.

Frisch, K. von and M. Lindauer (1954). “Himmel und Erde in Konkurrenz bei der Orientierung der Bienen”. In: Naturwiss 41, pp. 245–253.

Furlan, A., T. Baldwin, and A. Klippel (2007). “Landmark classification for route directions”. In: Proceedings of the Fourth ACL-SIGSEM Workshop on Prepositions. Association for Computational Linguistics. Prague, Czech Republic, pp. 9–16.

Geva-Sagiv, M., L. Las, Y. Yovel, and N. Ulanovsky (2015). “Spatial cognition in bats and rats from sensory acquisition to multiscale maps and navigation”. In: Nat Rev. Neurosci 16.2, pp. 94–108.

Giurfa, M., B. Eichmann, and R. Menzel (1995). “Symmetry as a perceptual category in honeybee vision”. In: Learning and memory. Proceedings of the 23rd Göttingen Neurobiology Conference. Ed. by N. Elsner and R. Menzel. Stuttgart: Thieme Verlag, p. 423.

Giurfa, M. (2003). “Cognitive neuroethology: dissecting non-elemental learning in a honeybee brain”. In: Current Opinion in Neurobiology 13.6, pp. 726–735.

Giurfa, M. (2015). “Learning and cognition in insects”. In: WIREs Cognitive Science 6.4, pp. 383–395.

Giurfa, M., S. Zhang, A. Jenett, and R. M. andMandyam V. Srinivasan (2001). “The concepts of ‘sameness’ and ‘difference’ in an insect”. In: Nature 410, pp. 930–933.

Gregory, R. L. and O. L. Zangwill, eds. (1987). The Oxford Companion to the Mind. Oxford University Press.

Heithaus, E. R., T. H. Fleming, and P. A. Opler (1975). “Foraging patterns and resource utilization in seven species of bats in a seasonal tropical forest”. In: Ecology 56, pp. 841–854.

Hentschke, H. and M. C. Stüttgen (2011). “Computation of measures of effect size for neuroscience data sets”. In: European Journal of Neuroscience 34.12. Toolbox available at https://github.com/hhentschke/measures-of-effect-size-toolbox, pp. 1887–1894.

Jacobs, L. F. and R. Menzel (2014). “Navigation outside of the box: what the lab can learn from the field and what the field can learn from the lab”. In: Movement Ecology 2.1, pp. 1–22.

Kehoe, E. J. (2008). “Discrimination and generalization”. In: Learning and Memory: A Comprehensive Reference. Ed. by J. H. Byrne and R. Menzel. Oxford: Academic Press. Chap. 1.08, pp. 123–149.

Kruskal, W. H. and W. A. Wallis (1952). “Use of ranks in one-criterion variance analysis”. In: Journal of the American Statistical Association 47.260, pp. 583–621.

Kurzhanskiy, A. A. and P. Varaiya (2006). “Ellipsoidal toolbox (ET)”. In: Proceedings of the 45th IEEE Conference on Decision and Control, pp. 1498–1503.

Landler, L., G. D. Ruxton, and E. P. Malkemper (2019). “Circular statistics meets practical limitations: a simulation-based Rao’s spacing test for non-continuous data”. In: Movement Ecology 7.1, 15.

Lipp, H.-P. et al. (2004). “Pigeon homing along highways and exits”. In: Current Biology 14.14, pp. 1239–1249.

Löfberg, J. (2004). “YALMIP: a toolbox for modeling and optimization in MATLAB”. In: 2004 IEEE International Conference on Robotics and Automation, pp. 284–289.

Mardia, K. V. (1972). Statistics of Directional Data. Probability & Mathematical Statistics Monograph. Academic Press.

McGill, R., J. W. Tukey, and W. A. Larsen (1978). “Variations of box plots”. In: The American Statistician 32.1, pp. 12–16.

McNamara, T. P., B. Rump, and S. Werner (2003). “Egocentric and geocentric frames of reference in memory of large-scale space”. In: Psychonomic Bulletin & Review 10.3, pp. 589–595.

Menzel, R., R. Brandt, A. Gumbert, B. Komischke, and J. Kunze (2000). “Two spatial memories for honeybee navigation”. In: Proceedings of the Royal Society of London. Series B: Biological Sciences 267.1447, pp. 961–968.

Menzel, R., K. Geiger, L. Chittka, et al. (1996). “The knowledge base of bee navigation”. In: Journal of Experimental Biology 199.1, pp. 141–146.

Menzel, R., K. Geiger, J. Joerges, U. Müller, and L. Chittka (1998). “Bees travel novel homeward routes by integrating separately acquired vector memories”. In: Animal Behaviour 55.1, pp. 139–152.

Menzel, R. and U. Greggers (2015). “The memory structure of navigation in honeybees”. In: Journal of Comparative Physiology A 201.6, pp. 547–561.

Menzel, R., U. Greggers, et al. (2005). “Honey bees navigate according to a map-like spatial memory”. In: Proceedings of the National Academy of Sciences 102.8, pp. 3040–3045.

Menzel, R., A. Kirbach, et al. (2011). “A common frame of reference for learned and communicated vectors in honeybee navigation”. In: Current Biology 21.8, pp. 645–650.

Menzel, R., L. Tison, et al. (2019). “Guidance of navigating honeybees by learned elongated ground structures”. In: Frontiers in Behavioral Neuroscience 12.

Mises, R. von (1918). “Über die ‘Ganzzahligkeit’ der Atomgewichte und verwandte Fragen”. In: Physikalische Zeitschrift 19, pp. 490–500.

Rescorla, R. A. (1992). “Response-outcome versus outcome-response associations in instrumental learning”. In: Animal Learning & Behavior 20, pp. 223–232.

Ruxton, G. D. (2017). “Testing for departure from uniformity and estimating mean direction for circular data”. In: Biology Letters 13.1, 20160756.

Schnute, J. T. and K. Groot (1992). “Statistical analysis of animal orientation data”. In: Animal Behaviour 43.1, pp. 15–33.

Se, S., D. G. Lowe, and J. J. Little (2005). “Vision-based global localization and mapping for mobile robots”. In: IEEE Transactions on Robotics 21.3, pp. 364–375.

Steel, E. A., M. C. Kennedy, P. G. Cunningham, and J. S. Stanovick (2013). “Applied statistics in ecology: common pitfalls and simple solutions”. In: Ecosphere 4.9, 115.

Stürzl, W., J. Zeil, N. Boeddeker, and J. M. Hemmi (2016). “How wasps acquire and use views for homing”. In: Current Biology 26.4, pp. 470–482.

Wehner, R. and R. Menzel (1990). “Do insects have cognitive maps?” In: Annual Review of Neuroscience 13.1, pp. 403–414.

Wehner, R., B. Michel, and P. Antonsen (1996). “Visual navigation in insects: coupling of egocentric and geocentric information”. In: Journal of Experimental Biology 199.1, pp. 129–140.

Wehner, R. and M. V. Srinivasan (1981). “Searching behaviour of desert ants, genus *Cataglyphis* (formicidae, hymenoptera)”. In: Journal of Comparative Physiology 142, pp. 315–338.

Wiener, J. et al. (2011). “Animal navigation: a synthesis”. In: Animal Thinking: Contemporary Issues in Comparative Cognition. Ed. by R. Menzel and J. Fischer. Strüngmann Forum Reports. Cambridge/USA: MIT Press, pp. 51–76.

Zentall, T. R., E. A. Wasserman, O. F. Lazareva, R. K. R. Thompson, and M. J. Rattermann (2008). “Concept learning in animals”. In: Comparative Cognition & Behavior Reviews 3, pp. 13–45.

